# Incongruence between transcriptional and vascular pathophysiological cell states

**DOI:** 10.1101/2023.04.16.537057

**Authors:** Macarena Fernández-Chacón, Severin Mühleder, Álvaro Regano, Lourdes Garcia-Ortega, Carlos Torroja, Mariya Lytvyn, Maria S Sanchez-Muñoz, Verónica Casquero-Garcia, Emilio Camafeita, Jesus Vazquez, Alberto Benguría, Ana Dopazo, Fátima Sánchez-Cabo, Hannah Carter, Rui Benedito

## Abstract

The Notch pathway is a major regulator of transcriptional specification and vascular biology. Previous studies have suggested that targeting the ligand Dll4 or the Notch-receptors results in similar molecular and angiogenesis outcomes. Here, we analyzed single and compound genetic mutants for all Notch signaling members and found very significant differences in the way ligands and receptors regulate vascular homeostasis. Loss of Notch receptors, leads to minor vascular pathology featuring hypermitogenic MAPK-driven cell-cycle arrest and senescence. In contrast, loss of Dll4 triggers a strong Myc-driven switch towards cell proliferation and sprouting and major organ pathology. Targeting of Myc completely suppressed the proliferative and tip-cell angiogenic states induced by Dll4 loss-of-function, however, this did not avoid vascular pathology. Only VEGF blockade prevented the pathology induced by Dll4 loss, but without fully suppressing its transcriptional and metabolic programs. This study shows incongruence between single-cell transcriptional states and adult vascular phenotypes and related pathophysiology.

## Introduction

Notch is a cell-to-cell ligand-receptor signaling pathway that has a major influence on cell transcription and biology^1,2^ playing important roles in several diseases^3^. γ-secretase inhibitors have been used to treat Alzheimer’s disease and cancer^4,5^. However, these general Notch signaling inhibitors have undesired effects, including disruption of the normal intestinal stem-cell differentiation^6,7^. Specific blocking antibodies are now available that target the various ligands and receptors of the Notch pathway ^8–15^. Given the specificity of Dll4 expression in endothelial cells, targeting this ligand was initially thought to be an effective and safe strategy for specifically modulating angiogenesis in disease, such as during tumor growth or tissue ischemia^12,13,16^. However, targeting Dll4 was later shown to induce a loss of endothelial quiescence and vascular neoplasms, causing major pathology in several organs ^10,14,17,18^.

Here, we characterized the impact on vascular quiescence of single or compound targeting of all Notch signaling members. High-resolution scRNAseq and 3D confocal microscopy revealed highly significant differences in the way each Notch member regulates vascular signaling, homeostasis, and single-cell states in diverse vascular beds. γ-secretase inhibitors or removal of Notch receptors did not cause significant vascular or organ disease. Abnormal proliferating and sprouting single-cell states were generated only by continuous targeting of Dll4. Surprisingly, suppression of these abnormal cell states by additional genetic or pharmacological targeting was insufficient to prevent vascular and organ disease. Our results show that the transcriptional changes and angiogenic cell states elicited by anti-Dll4 are not what causes vascular pathophysiology. Instead, we propose that is the unrelated vascular enlargement and malfunction that leads to organ pathology.

## Results

### Expression and function of Notch signaling members in distinct organ vessels

To elucidate the role of Notch signaling in global vascular homeostasis, we first assessed its activity in different organ vascular beds by immunodetection of the activated form of the Notch1 intracellular domain (N1ICD^Val1744^)^2^. This epitope was detected in ~50% of all organ ECs (Fig. 1a, 1b). Bulk RNAseq analysis revealed that *Dll4* and *Notch1* are the most expressed ligand–receptor pair in quiescent vessels of most organs (Fig. 1c, 1d, Supplementary Fig. 1a), and that *Mfng* is the most strongly expressed Notch glycosyltransferase. These enzymes are known to significantly enhance Delta ligand signaling and decrease Jagged ligand signaling^19,20^. *Dll4* gene deletion in adult *Dll4^flox/flox^ Cdh5-CreERT2* mice (abbreviated as *Dll4^iDEC^*, with *Dll4* induced deletion in ECs) led to a significant reduction in N1ICD^Val1744^ and Hey1 signals in quiescent ECs of *Dll4^iDEC^* organs (Fig. 1e-i). This indicates that Dll4 is the main functional ligand responsible for triggering Notch activity in most quiescent vessels. Only in Lungs we observed compensatory upregulation of *Dll1* (Fig. 1i). *Dll4* deletion elicited remarkably different gene expression signatures among different organ vascular beds, with the adult liver endothelium presenting the most pronounced changes in gene expression (Fig. 1j, k and Supplementary Fig. 1). Despite significant transcriptional changes in most organs endothelial cells, only the endothelium of the heart, muscle, and liver showed an increase in the frequency of cycling or activated Ki67+ cells upon *Dll4* deletion (Fig. 1l-n), and these were the only organs with clear alterations in the 3D vascular architecture after the loss of Dll4– Notch signaling (Fig. 1o). The brain underwent significant changes in gene expression (Fig. 1j, k and Supplementary Fig. 1), but these were not accompanied by endothelial proliferation or vascular morphological changes.

**Figure 1:**
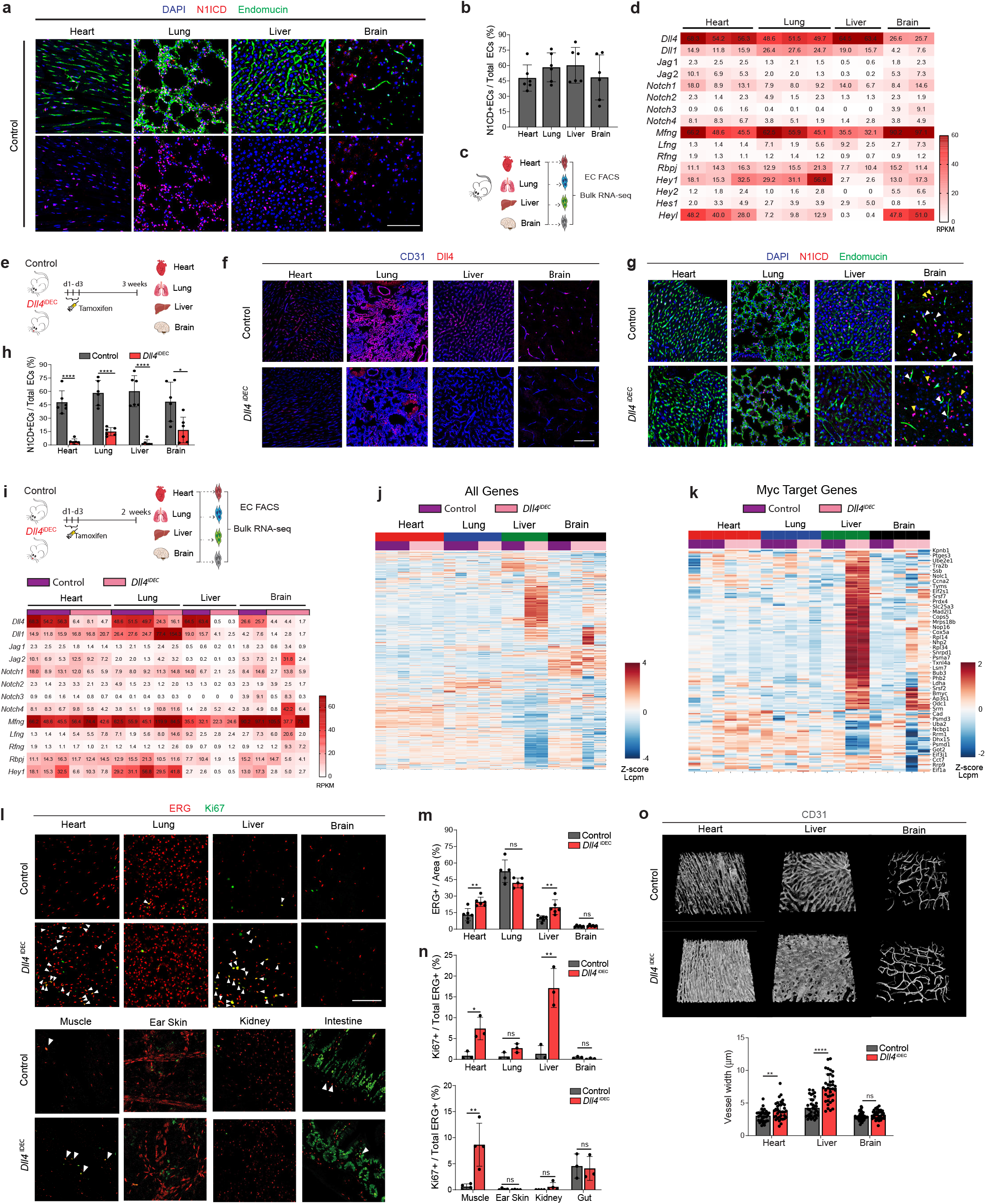
*Dll4* deletion leads to EC activation and proliferation only in some vascular beds. **a,b,** Notch1 signalling activity (cleaved Val1744 N1ICD) in quiescent endothelium (DAPI+Endomucin+). **c,** Schematic representation to illustrate the bulk RNA-seq experiment performed with adult ECs isolated by FACS. **d,** Heatmap with RNA-seq RPKM (Reads per kilo base per million mapped reads). **e,** Experimental layout for the inducible deletion of *Dll4* in Cdh5+ ECs (*Dll4*^iDEC^) with *Cdh5(PAC)-CreERT2*. **f,** Expression of Dll4 protein in CD31+ vessels. **g,** *Dll4* deletion significantly reduces Notch signalling activity (cleaved Val1744 N1ICD) in all quiescent vascular beds. In brain micrographs, white arrowheads indicate ECs and yellow arrowheads non-ECs. Note that whereas N1ICD is maintained in non-ECs, most N1ICD signal disappears from the ECs in *Dll4^iDEC^* brains. **i,** Schematic representation to illustrate the bulk RNA-seq experiment performed with adult ECs. Below a heatmap showing the relative expression of all Notch pathway components and canonical target genes in control and *Dll4*^iDEC^ endothelial cells. **j,** Unsupervised hierarchical clustering showing stronger gene expression changes in *Dll4*^iDEC^ liver ECs compared to the other organs. Z-score lcpm: Z-score of the logarithmic counts per million. **k,** Unsupervised hierarchical clustering showing strong upregulation of Myc targets genes in *Dll4*^iDEC^liver ECs compared to the other organs. **l-n,** Dll4 deletion results in increased EC proliferation (Ki67+ERG+ cells) in some organs but not others. **o,** 3D reconstruction images from thick vibratome sections show vessel (CD31+) enlargement in *Dll4*^iDEC^ heart and liver but not in brain. Charts n=3 animals per group minimum. Error bars indicate SD. *p < 0.05. **p < 0.01. ****p < 0.0001. ns, non-significant. Scale bars, 100 μm.

### Targeting Dll4 induces heterozonal responses in liver vessels

The RNAseq and histological data revealed the adult liver endothelium as the most reactive vascular bed to the targeting of Dll4–Notch signaling. Rodents and chimpanzees treated with anti-Dll4 antibodies develop significant liver disease, and because this was suggested to be one of the main barriers to the therapeutic use of anti-Dll4 or other Notch inhibitors^10,14^, we focused our analysis on this organ. To gain deeper insight, we performed a high-resolution spatiotemporal phenotypic and transcriptomic analysis after targeting Dll4 for 2 days to 3 weeks. In contrast to targeting Dll4 during angiogenesis, targeting Dll4 in quiescent liver vessels for 48h, which abolishes the generation of cleaved N1ICD, did not induce major transcriptomic changes (only 11 DEGs) or vascular phenotypic changes (Supplementary Fig. 2a-e). Gene-set enrichment analysis (GSEA) revealed upregulation of only a few E2F and Myc target genes at this time point (Supplementary Fig. 2f-2h). The increase in vascular density was relatively slow and progressive, only becoming noticeable more than a week after *Dll4* deletion (Fig. 2a-c). This was associated with a peak in endothelial proliferation rate at day 4, which was sustained until at least the 3rd week, leading to a progressive increase in vascular density and the total number of ECs (Fig. 2b-d). This endothelial activation induced by Dll4 targeting also induced increased proliferation of neighboring hepatocytes, peaking after the peak in endothelial proliferation (Fig. 2e) and suggesting Dll4^KO^ ECs secrete angiocrine factors inducing hepatocyte proliferation, as shown before during liver regeneration^21,22^.

**Figure 2:**
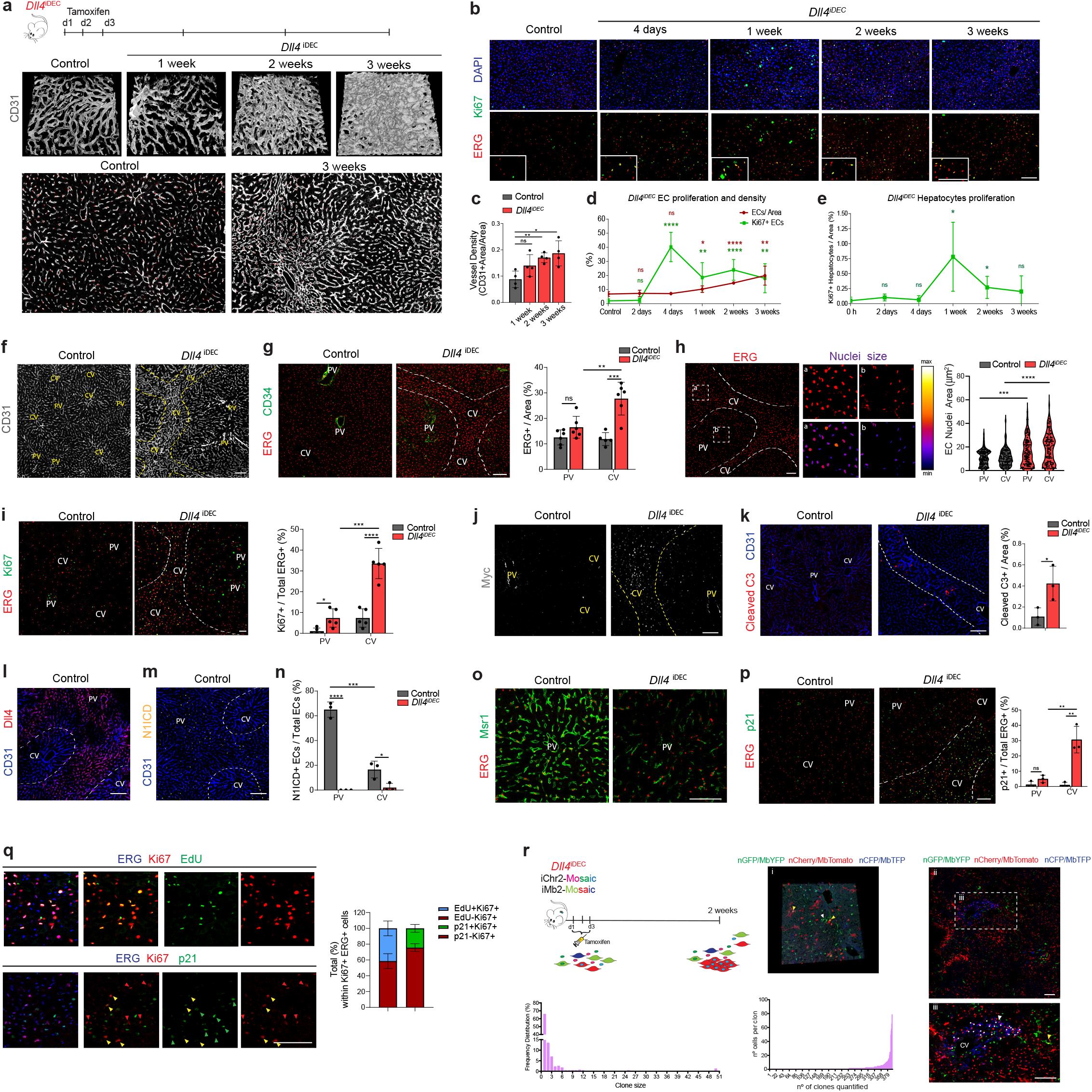
Targeting Dll4 induces heterozonal responses in liver vessels. **a,** Experimental layout for the inducible deletion of *Dll4* in Cdh5+ ECs (*Dll4*^iDEC^) with *Cdh5(PAC)-CreERT2*. 3D projection of confocal images from thick vibratome sections show that vascular malformations and capillary enlargement arise after the first week post-*Dll4*-deletion. **b-e,** Analysis of EC (ERG+ cells) and hepatocyte (ERG-DAPI+) proliferation (Ki67+) and cell number over time. **f**, Representative confocal micrographs showing that the enlarged and abnormal vascular pattern observed in *Dll4*^iDEC^ livers is located in the central veins (CV)-connecting sinusoids, but not in ECs surrounding portal veins (PV). Yellow dashed lines highlight the CV affected area. **g**, EC density in *Dll4*^iDEC^ liver is higher in the endothelium connecting the CVs rather than the endothelium surrounding PVs, that are distinguished by CD34+ staining. White dashed lines highlight the denser area. **h,** *Dll4*^iDEC^ liver section showing the increase in nuclei size mainly in CV-connecting sinusoids. White dashed lines highlight the area with higher EC density and with larger EC nuclei. Higher magnification pictures of insets (a) and (b) together with pseudocolouring of nuclear sizes (lower panels) show differences in nuclei size between CV and PV areas, respectively. Violin plots reflecting changes in cell nuclei sizes. **i**, Increased EC proliferation (Ki67+ERG+) in *Dll4*^iDEC^ liver particularly in the sinusoids connecting the CVs. **j**, Myc protein is upregulated mainly in ECs (ERG+ cells) around the CVs after *Dll4* deletion. **k**, Increased apoptosis (cleaved Caspase 3) in CV areas upon *Dll4* deletion. **l-m**, Dll4 and activated N1ICD (V1744) protein are mostly present in arterial PV areas, while being mostly undetectable in venous CV areas. **n,** *Dll4* deletion leads to loss of N1ICD (Val1744) activation in liver ECs. **o**, Msr1 immunostaining showing loss of arterial identity in *Dll4*^iDEC^ vessels. **p,** p21 expression in *Dll4*^iDEC^ liver ECs (ERG+) is also higher in the sinusoids around the CVs. **q,** *Dll4*^iDEC^ Ki67+ liver ECs are actively dividing in S-phase (EdU+Ki67+ERG+) and a small fraction of proliferating ECs (Ki67+ERG+) also expresses p21 protein (p21+Ki67+ERG). **r,** Dual ifgMosaic tracking reveals that few ECs (<25%) have the ability to divide and clonally expand after Dll4 targeting. Images showing representative dual-labelled EC clones (yellow and white arrowheads in (i) and asterisks in (iii)) showing that proliferative ECs are located in endothelial sinusoids around the CVs. Charts n=3-4 animals per group minimum. Error bars indicate SD. *p < 0.05. **p < 0.01. ***p < 0.001. ****p < 0.0001. ns, non-significant. Scale bars, 100 μm.

The effect of Dll4 targeting was, however, notably heterogeneous and zonal. Only vessels around the central veins (CVs) and with a known venous identity^23,24^ showed an abnormal vascular pattern, featuring a higher number of ECs (Fig. 2f, g), larger nuclei (Fig. 2h), and expression of cell cycle (Fig. 2i, 2j) and apoptosis (Fig. 2k) markers. We also observed that the previously reported anti-Dll4–driven liver histopathology^14,18^ was exclusively associated to the abnormal central-vein sinusoids (Supplementary Fig. 2i-2l). Paradoxically, the portal vein (PV) vessels, which have arterial identity and the highest Dll4 expression and Notch activity (Fig. 2l-n), conserved normal vascular architecture (Supplementary Fig. 2i-l) and showed only a very subtle increase in EC proliferation (Fig. 2i). Normal arterial vascular architecture and function was maintained despite the lack of Notch signaling and arterial identity in *Dll4^iDEC^* vessels (Fig. 2n, o and Supplementary Fig. 2m). Besides the cell-cycle marker Ki67, we also analyzed more specific S-phase (EdU) and cell-cycle arrest/senescence (p21) markers. This analysis revealed expression of p21 in 30% of *Dll4^iDEC^* ECs in the venous vessels around the central veins (Fig. 2p). Among Ki67+ ECs, 40% were positive for EdU and 25% for p21 (Fig.2q). This shows that there is a mix of productive cell division (EdU+) and arrest (p21+) after Dll4 loss in liver ECs. Pulse–chase single-cell ifgMosaic tracking revealed that despite 34% of all venous ECs being Ki67+, few of these ECs had the ability to divide and clonally expand after Dll4 targeting, with some cells dividing 6 to 50 times more than their neighbors (Fig. 2r histograms). All of these resident progenitor cells were located in the sinusoids around central veins (Fig. 2r iii).

### Loss of Notch1 or Rbpj leads to non-pathologic hypermitogenic cell-cycle arrest

Notch ligands and receptors can be targeted with a range of pharmacological compounds and antibodies^8-13^, and so far only Dll4-targeting antibodies have been found to cause major vascular disease^10,25^. In contrast, genetic deletion of *Notch1* or *Rbpj* in mice has been suggested to cause vascular phenotypes very similar to the genetic deletion of *Dll4*, during angiogenesis and in adult vessels^17,26–28^. We therefore investigated if deleting *Notch1* or *Rbpj,* the master regulator of all Notch receptor signaling, induced vascular pathology similar to that induced by the loss of Dll4 (Fig. 3a). Surprisingly, *Notch1* and *Rbpj* deletion did not significantly increase EC proliferation or induce liver disease (Fig. 3b-e, Supplementary Fig. 3a, b), despite these mutant cells having even higher phospho-ERK (P-ERK) activity than ECs lacking Dll4 (Fig. 3f-h). We next compared the transcriptome of *Dll4^iDEC^* and *Rbpj^iDEC^* vessels. ECs from both mutant lines showed upregulation of genes related to cell-cycle activation and metabolism (Fig. 3i) and had enlarged nuclei (Fig. 3j). However, compared with *Dll4^iDEC^* livers, *Rbpj^iDEC^*livers had less vascular dilation and organ abnormalities (Fig. 3k, Supplementary Fig. 3b) and stronger upregulation of p21 (Fig. 3l), a cell-cycle inhibitor frequently upregulated in senescent or hypermitogenically arrested cells^29^. We also identified a significant increase in the number of binucleated p21+ ECs, suggestive of replicative stress and G2 arrest of the mutant cells (Fig. 3m and Supplementary Fig. 3c). RNaseq analysis also revealed signatures of genetic pathways linked to p53, p21, G2M checkpoints, chromosome segregation, and general replicative stress and senescence in *Rbpj^iDEC^*ECs (Fig. 3n, Supplementary Fig. 3d). To determine the functional impact of p21 upregulation, we analyzed compound *Rbpj^iDEC^ p21^KO^* mice (Fig. 3o). p21 loss did not affect the minor vascular sinusoid dilation seen in *Rbpj^iDEC^* livers, but did increase the frequency of cycling (Ki67+) and apoptotic (Cleaved caspase3+) cells (Fig. 3p-r), in line with the role of p21 as a cell-cycle and apoptosis inhibitor^30,31^, particularly in hypermitogenically activated *Rbpj^KO^* cells. This dual and paradoxical effect of p21 loss on both cell proliferation and apoptosis may explain the relatively mild increase in EC numbers in *Rbpj^iDEC^ p21^KO^* livers compared with the fully arrested *Rbpj^iDEC^* liver vessels. These results suggest that loss of Dll4 induces a reduction in Notch signaling that results in a mixed population of proliferative and arrested ECs, whereas the complete loss of Notch signaling in *Notch1^iDEC^* and *Rbpj^iDEC^* mice induces mostly hypermitogenic arrest, without cell division.

**Figure 3:**
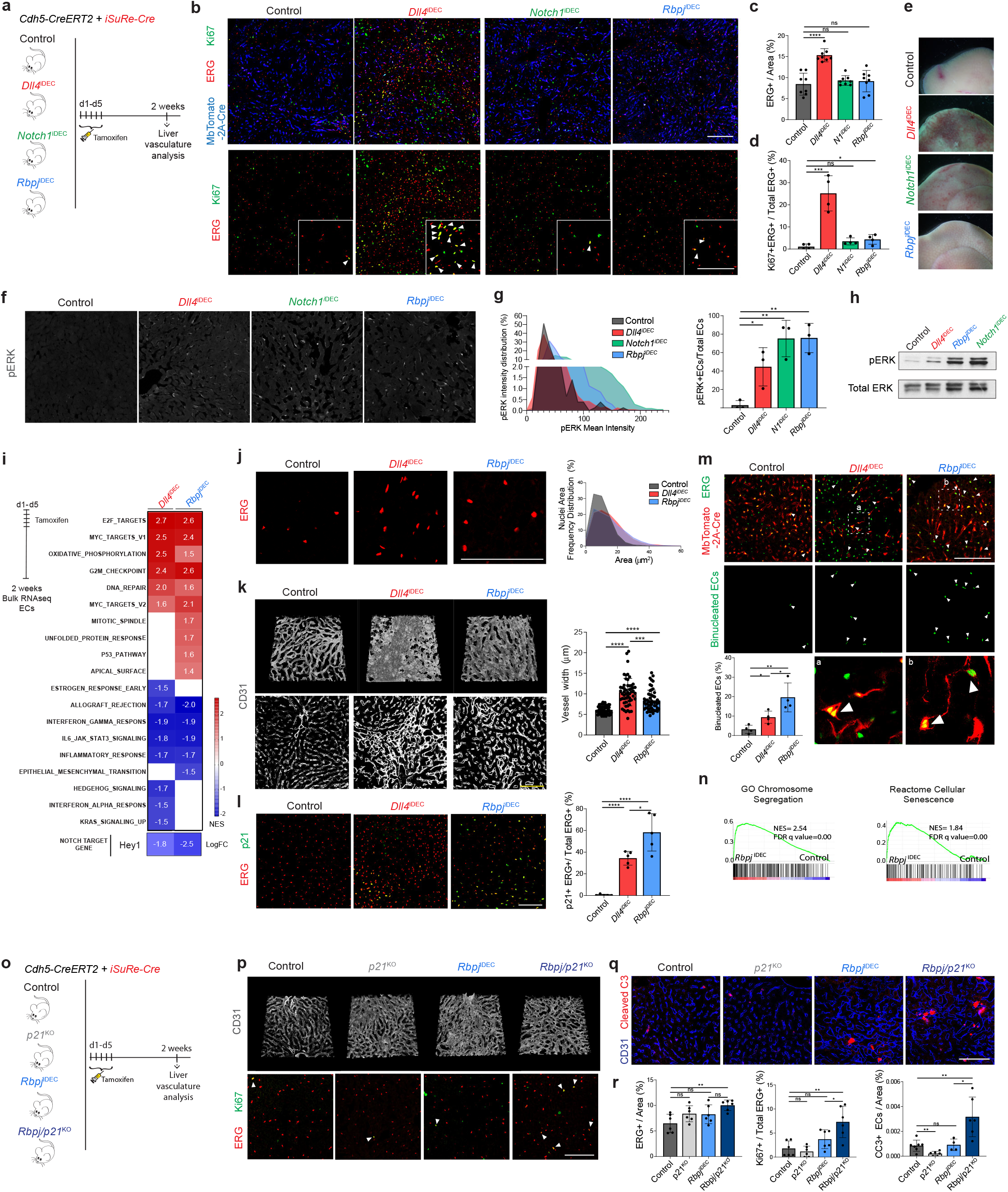
Deletion of *Rbpj* or *Notch1* in liver quiescent blood vessels does not phenocopy *Dll4* deletion. **a,** Experimental layout for the inducible deletion of *Rbpj* (*Rbpj*^iDEC^), *Notch1(Notch1*^iDEC^) and *Dll4*(*Dll4*^iDEC^) in Cdh5+ ECs. All mice contained the *Cdh5(PAC)-CreERT2* and *iSuRe-Cre* (expressing MbToma-to-2A-Cre) alleles to ensure genetic deletion of the floxed alleles. **b-d,** Increased EC density (ERG+ per field) and proliferation (Ki67+ERG+/ERG+) was observed only in *Dll4^i^*^DEC^ liver ECs. **e,** Gross liver pathology is observed exclusively in *Dll4^iDEC^* livers. **f-h,** p-ERK immunostaining and whole liver western blot showing that the frequency of p-ERK expressing ECs and intensity levels increase in the mutants, particularly the Notch1 and Rbpj mutants. **i,** Heatmap with the normalized enrichment score (NES) from significant Hallmark analysis (FDR qval< 0.05) by GSEA from bulk RNAseq data. **j,** Mutant liver ECs have a larger nuclei size compared with control liver ECs. **k,** Vascular (CD31+) dilation/expansion is more pronounced in *Dll4*^iDEC^ mutants. **l,** p21 expression in ECs (p21+ERG+) is more increased in *Rbpj*^iDEC^ mutants. **m,** Binucleated cells (white arrowheads) identified in *Dll4*^iDEC^ and *Rbpj*^iDEC^ mutants. High magnification of insets (a) and (b) are shown at the bottom. **n,** GSEA analysis show a positive and significant enrichment in Chromosome Segregation and Cellular Senescence-related genes in *Rbpj*^iDEC^ mutant liver ECs as shown by the Normalized Enrichment Score (NES). **o,** Experimental layout for the inducible deletion of *Rbpj* in a *p21^KO^* background. **p,** 3D projection of thick vibratome sections showing the endothelial surface marker CD31, and proliferation (Ki67) analysis in ECs (ERG+). **q,** Analysis of the apoptosis marker cleaved caspase 3. **r,**The absence of p21 in a *Rbpj*^iDEC^ background results in a modest increase in EC density (ERG+), but both EC proliferation (Ki67+ERG+) and apoptosis (cleaved Caspase 3) are significantly increased. Charts n=3-5 animals per group minimum. Error bars indicate SD. *p < 0.05. **p < 0.01. ***p < 0.001. ****p < 0.0001. ns, non-significant. Scale bars, 100 μm.

### Targeting Dll4 and Notch receptors induces incongruent cell states

We next performed single cell RNAseq to identify possible differences in vascular single-cell states induced by targeting Dll4, Notch1, and Rbpj. This analysis was performed on cells expressing the *Cdh5-CreERT2* and *iSuRe-Cre* alleles^32^ to guarantee endothelium-specific recombination, labeling, and full genetic deletion of all the floxed genes used in this study (Fig. 4a-b, Supplementary Fig. 4a). To reduce batch effects, Tomato+CD31+ endothelial cells were isolated on the same day from multiple control and mutant animals, tagged with different oligonucleotide-conjugated antibodies, and loaded in the same chip. The few mutant cells with mRNA expression of *Dll4* and *Notch1* were likely fluorescence-activated cell sorting (FACS) contaminants. In the case of *Rbpj*, only exons 6-7 are deleted, leading to a less stable, but still detectable, 3′ mRNA. Altogether, the scRNAseq data analysis showed the existence of 10 clearly defined cell clusters (Fig. 4c-e, Supplementary Fig. 4b). The deletion of *Rbpj, Notch1*, and *Dll4* resulted in a significant decrease in Notch signaling and Hes1 expression (Fig. 4b) and in the loss of the arterial sinusoidal capillaries C1a cluster; however, only the loss of Dll4 was able to induce a very pronounced tip-cell transcriptional program (C4)(Fig. 4c-e). This program was characterized by the loss of venous *Wnt2* gene expression and very high expression of the tip-cell markers *Kcne3, Esm1, Angpt2* and *Apln,* as well as *Myc* and its canonical target *Odc1* (Fig. 4e,f and Supplementary Fig. 4b, 4c). Most of the upregulated genes in the tip-cell cluster were associated with Myc metabolism, increased ribosome biosynthesis, glycolysis, mTorc1 signaling, and also fatty acid and oxidative phosphorylation (Supplementary Fig. 4d). Paradoxically, *Notch1^iDEC^* and *Rbpj^iDEC^* liver ECs, in which the decrease in Notch signaling was more pronounced (*Hes1* expression in Fig. 4b), showed a more moderate metabolic activation, and most of these mutants ECs clustered either in the venous C1v or in the activated C3 cluster and did not reach the extreme C4 tip-cell state (Fig. 4c,d).

**Figure 4:**
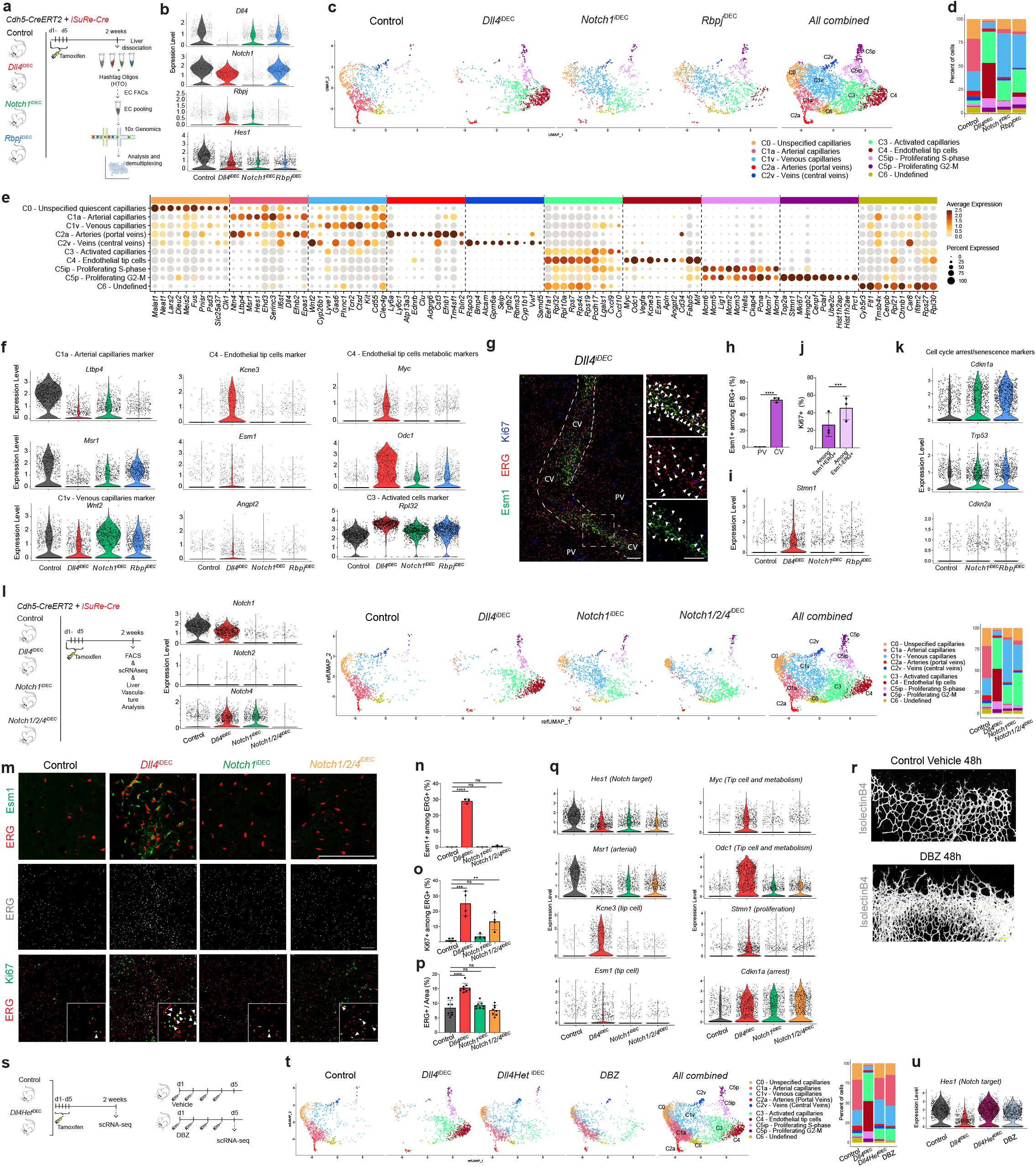
scRNAseq analysis reveals significant differences between targeting Dll4 and Notch signalling. **a,** Experimental layout for the inducible deletion of the indicated genes in Cdh5-CreERT2+ ECs and collection of the iSuRe-Cre+ (Tomato-2A-Cre+) cells to ensure genetic deletion. **b,** Violin plots showing *Dll4*, *Notch1* and *Rbpj* mRNA expression in single cells and the subsequent downregulation of the Notch target gene *Hes1* in all mutants. **c,** UMAPs showing the 10 identified cell clusters and subclusters in the scRNAseq. **d,** Barplot showing the percentage of cells from each cluster in the different samples. **e,** Dot plot showing the frequency (size) and intensity (color) of expression for the top cluster marker genes. **f,** Violin plots for some cluster marker genes. **g-h,** In *Dll4*^iDEC^ mutants, cells from the tip cell C4 cluster (Esm1+ERG+) are localized in the sinusoids around central veins (CVs), but not in PVs. **i,** The global cell-cycle marker *Stmn1* is highly upregulated exclusively in *Dll4*^iDEC^ liver ECs. **j,** Most of the Esm1+ECs are not Ki67+, but have Esm1-Ki67+ECs+ as neighbours in the CV sinusoids. **k,** Violin plots showing upregulation of Cdkn1a (p21), Trp53 (p53) and Cdkn2a (p16) in *Notch1*^iDEC^ and *Rbpj*^iDEC^ mutant ECs. **l,** Experimental layout for the inducible deletion of the indicated genes in Cdh5-CreERT2+ ECs and collection of the iSuRe-Cre+ (Tomato-2A-Cre+) cells to ensure genetic deletion. Violin plots showing expression of the indicated genes in the isolated single ECs. UMAPs and barplot showing the 10 identified clusters and the percentage of cells for each cluster in the different samples. **m-p,** Full loss of Notch receptors does not induce the tip cell marker Esm1. It does induce an increase in Ki67+ ECs, but this does not result in increased EC numbers (ERG+). **q,** Violin plots confirming that deletion of *Notch1/2/4* does result in less Notch signalling (*Hes1*), and less arterial marker expression (*Msr1*), but this does not induce the tip cell programme (*Kcne3/Esm1/Myc/Odc1*) or the proliferation marker *Stmn1* observed in *Dll4*^iDEC^ mutants. The cell-cycle arrest marker *Cdkn1a*/p21 is highly expressed. **r,** DBZ (Y-secretase/Notch inhibitor) treatment for 48h in developing retina vessels leads to increased vessel density, as observed after anti-Dll4 treatment. **s,** Experimental layout for the inducible heterozygous deletion of *Dll4* (*Dll4Het*^iDEC^) for 2 weeks or DBZ treatment for 4 days in Cdh5+ ECs and scRNAseq data comparison. **t,** UMAPs and barplot show the 10 identified clusters and the percentage of cells in each cluster in the different samples. **u,** Violin plots showing that unlike *Dll4*^iDEC^, DBZ treatment and the heterozygous deletion of *Dll4* have a weak effect on *Hes1* expression. n=3-4 animals per group minimum. Error bars indicate SD **p < 0.01. ***p < 0.001. ****p < 0.0001. ns, non-significant.Scale bars, 100 μm.

Histology confirmed that indeed only the *Dll4^iDEC^* mutants had a significant population of Esm1+ tip cells (Supplementary Fig. 4e) and these were mostly present in the venous sinusoidal capillaries interconnecting the liver central veins (Fig. 4g, h), where EC proliferation and density are the highest (Fig. 2f-i). Our previous immunostaining analysis revealed that around 25% of Dll4 mutant cells were Ki67+, in contrast to 4% in Notch1 and Rbpj mutants and 1% in wildtype animals (Fig. 3b, d). The upregulation of the global cell-cycle marker *Stmn1* in *Dll4^iDEC^* livers correlated with the 6-fold higher frequency of Ki67-protein+ cells in these mutants than in the Notch1 and Rbpj mutants (Fig. 4i). Interestingly, most Esm1+ tip cells were not Ki67+, in accordance with their higher sprouting activity and arrested nature, but had proliferating Ki67+ cells as close neighbors (Fig. 4g, j). The scRNAseq data revealed that the number of activated cells (C3) and expressing S-G2-M markers (C5) was similar in the three mutants, even though Dll4 mutants had 6 times more Ki67+ and Stmn1+ cells. *Notch1^iDEC^* and *Rbpj^iDEC^* ECs also showed significant upregulation of the replication-stress/senescence markers p21 (*cdkn1a*), p53 (*trp53*), and p16 (*cdkn2a*) (Fig. 4k). These data suggests that many *Notch1^iDEC^* and *Rbpj^iDEC^*ECs undergo hypermitogenic S-G2-M arrest (Fig. 3m, 3n), without becoming Kcne3+/Esm1+ sprouting tip cells (Fig. 4c, d), which is in contrast to the current understanding of sprouting angiogenesis^33,34^.

*Notch1^iDEC^* livers upregulated the expression of *Notch4* (Supplementary Fig. 4f), a receptor known to partially compensate for *Notch1* deletion^35^. However, full loss of all Notch-receptor signaling in ECs did not induce the Esm1/Kcne3+ tip-cell state (Fig. 4l-n). Compared with *Notch1^iDEC^* livers, deletion of all receptors did induce a higher frequency of Ki67+ and activated/primed (C3) cells, which were nonetheless non-proliferative and non-pathologic and did not result in more ECs or increased vascular density (Fig. 4o, p, Supplementary Fig. 4g). In contrast to ECs losing Dll4, full loss of Notch receptors, similarly to Rbpj loss, results in even lower Hes1 expression and higher p21 expression (arrest); however, this does not result in the induction of tip cells (Esm1+/Kcne3+) or proliferating Stmn1+ cells (Fig. 4q).

We also tested a general γ-secretase inhibitor (DBZ) that is known to block Notch signaling and elicit strong effects on tumor and retina angiogenesis^12^, similarly to anti-Dll4 (compare Fig. 4r and Supplementary Fig. 2a). However, this compound had a very weak effect on quiescent vessels, inducing a minor increase in the C3 activated cluster and a minor decrease in the arterial capillary C1a cluster, similar to the changes seen in Dll4 heterozygous livers (Fig. 4s-u). We also observed by scRNAseq that ECs with full loss of Dll4 signaling for 4 days had already lost the arterial capillary program (C1a) and become activated (C3), but had not yet had time to fully differentiate to tip cells (C4 cluster in Supplementary Fig. 4h-j). This suggests that in order to fully activate quiescent ECs and induce significant numbers of tip cells and vascular abnormalization, pronounced and continuous loss of Dll4 signaling must be sustained for about a week, which can be achieved with genetic deletion or blocking antibodies^10^ but not with small molecule inhibitors targeting Notch.

### Deletion of all other Notch ligands leads to the non-pathologic upregulation of Notch signaling

Besides Dll4, other Notch ligands are also expressed in liver ECs (Fig. 5a). The Notch signaling target *Hes1* is more expressed in *Dll4^iDEC^* than in *Notch1^iDEC^*, *Rbpj^iDEC^*, or *Notch1/2/4^iDEC^* mutants (Fig. 4b, 4q), suggesting that the other weakly expressed Notch ligands (Dll1, Jagged1, and Jagged2) may partially compensate the loss of Dll4 and induce residual Notch signaling essential for the induction of the tip-cell state. Notably, *Jagged1* mRNA was barely detectable in bulkRNAseq data of quiescent livers (Fig. 1d), but its protein was strongly expressed in liver vessels (Fig. 5b). Deletion of all three ligands (*Jag1, Jag2,* and *Dll1*) did not alter vascular morphology, induce pathology, or increase the frequency of Ki67+ cells, confirming that Dll4 is the main Notch ligand in quiescent vessels (Fig. 5c-g). In agreement with this, scRNAseq data analysis confirmed that most cells remained quiescent and did not become activated or form tip cells (Fig. 5h-j). Moreover, unlike deletion of *Dll4*, *Notch1*, and *Rbpj*, triple *Jag1/2/Dll1* deletion in ECs did not compromise arterial identity (Fig. 5k), instead revealing a significant and counterintuitive increase in the Notch signaling target *Hes1* and an increase in the arterial gene *CD34*, together with a very pronounced decrease in the expression of the venous-enriched *Wnt2* gene (Figure 5l). This increased Notch signaling is also observed after the loss of Jagged ligands during angiogenesis^36^ and is likely related to the strong EC expression of Fringe glycosyltransferases (particularly *Mfng*, see Fig. 1d), previously shown to induce Delta ligand signaling while inhibiting Jagged ligands^19,20^, creating antagonistic and competitive signaling activity among distinct ligands binding the same Notch receptors. However, in the absence of the stronger Dll4 ligand, Jagged and Dll1 ligands can still induce residual Notch signaling (see Fig. 4b *Hes1*), that may be essential for effective induction of the endothelial tip-cell (C4) state and the observed vascular pathology.

**Figure 5:**
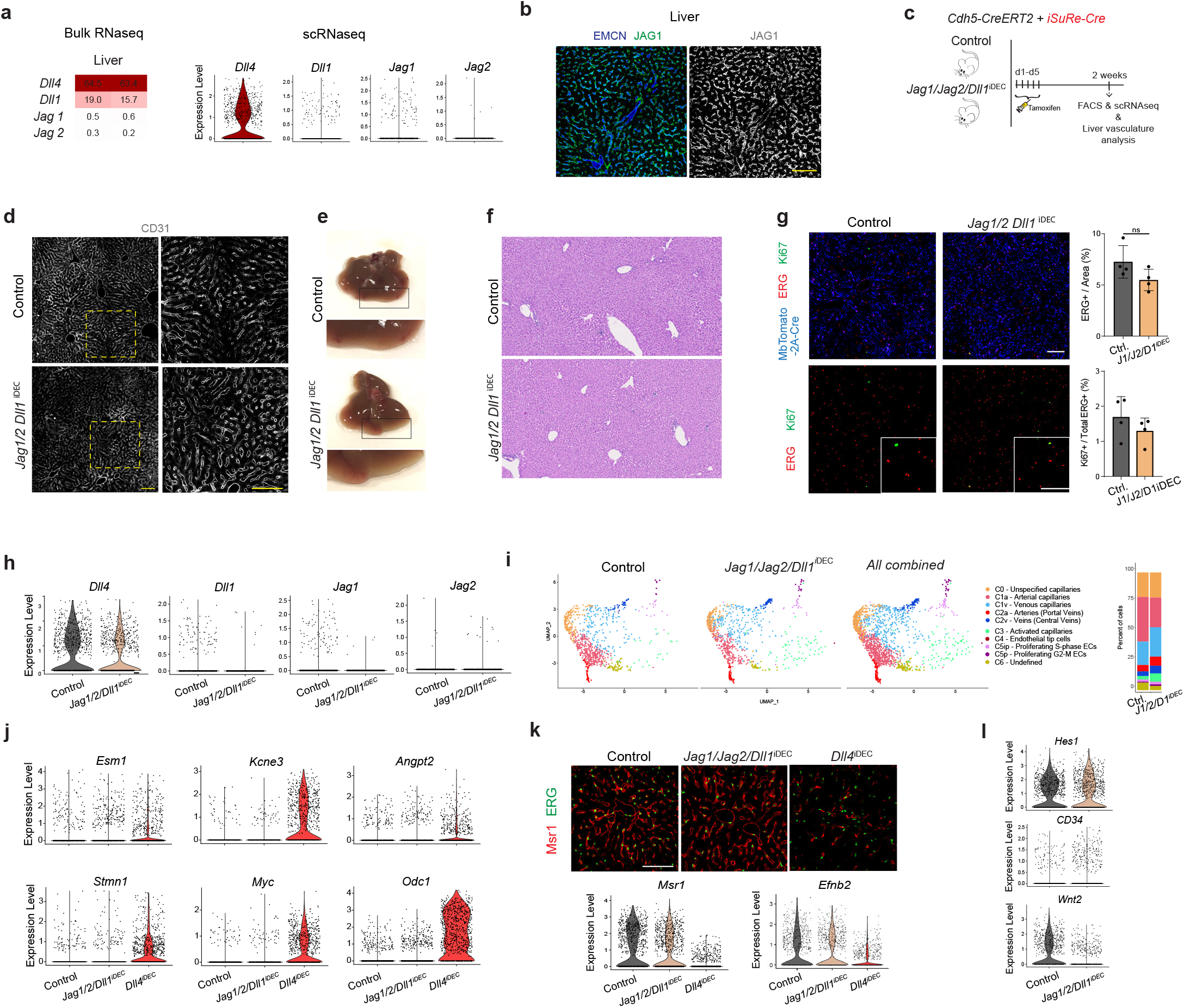
Deletion of *Jag1/Jag2/Dll1* lead to the non-pathologic upregulation of Notch signalling. **a,** Heatmap of bulk RNAseq data and violin plot of single cell data showing expression of all Notch ligands in liver ECs. **b,** Despite its low mRNA expression, Jag1 protein is strongly detected in the adult liver quiescent endothelium (CD31+). **c,** Experimental layout for the inducible deletion of *Jag1*, *Jag2* and *Dll1* in Cdh5+ ECs and histology or scRNAseq analysis. **d,** CD31+ immunostaining shows no vascular architecture changes. **e-f,** Macroscopic pictures and H&E staining show absence of liver pathology. **g,** Deletion of the three ligands does not lead to an increase in endothelial proliferation (Ki67+/ERG+ECs) or increase in EC number (ERG+ cells per field) compared to control livers. **h,** Violin plots showing expression of the 4 ligands in scRNAseq data. **i,** UMAPs and barplot showing the 10 identified clusters and the percentage of cells in each cluster in the two samples. **j,** Jag1/Jag2/Dll1 mutant ECs do not acquire the tip-cell (*Esm1/Kcne3/Angpt2*), nor metabolic (*Myc/Odc1*), nor proliferation (*Stmn1*) transcriptional programme observed in Dll4 mutants. **k,** Immunostaining and scRNAseq data showing that Jag1/Jag2/Dll1 mutant ECs do not downregulate the expression of the arterial markers *Msr1* and *Efnb2*. **l,** Violin plot showing an increase in the Notch target gene *Hes1* and the arterial gene *CD34*, together with a decrease in the expression of the venous *Wnt2* gene in Jag1/Jag2/Dll1 mutant ECs. Error bars indicate SD. ns, non-significant.Scale bar, 100 μm.

### Myc loss prevents the appearance of the *Dll4^KO^* single cell states but not vascular pathology

We next aimed to determine the molecular mechanisms responsible for the unique EC activation, tip-cell signature, and vascular pathology induced by targeting Dll4. All results so far led us to hypothesize that the vascular-related organ pathology in these mutants was caused by the high frequency of cells in the proliferative and tip-cell states. As mentioned above, *Myc* and its target *Odc1* were among the most strongly upregulated genes in Dll4 mutant ECs, compared with Notch1 and Rbpj mutants. Myc is known to activate important ribosome biogenesis and protein translation pathways, favoring cell growth^37^. *Dll4^iDEC^* livers showed upregulation of a large range of canonical E2F, Myc, and mTORC1 target genes and ribosomal (*Rpl*) genes, particularly in the activated, proliferating, and endothelial tip-cell clusters (Figure 6a, Supplementary Fig. 4). This hypermetabolic transcriptional status was confirmed by mass spectrometry (MS) analysis of protein lysates obtained from millions of freshly isolated liver ECs (Figure 6b-f), providing a first high-depth proteomic analysis of the endothelial tip-cell state induced by targeting Dll4. We also independently confirmed *Myc* mRNA and protein upregulation in *Dll4^KO^* vessels (Figure 6g-h).

**Figure 6:**
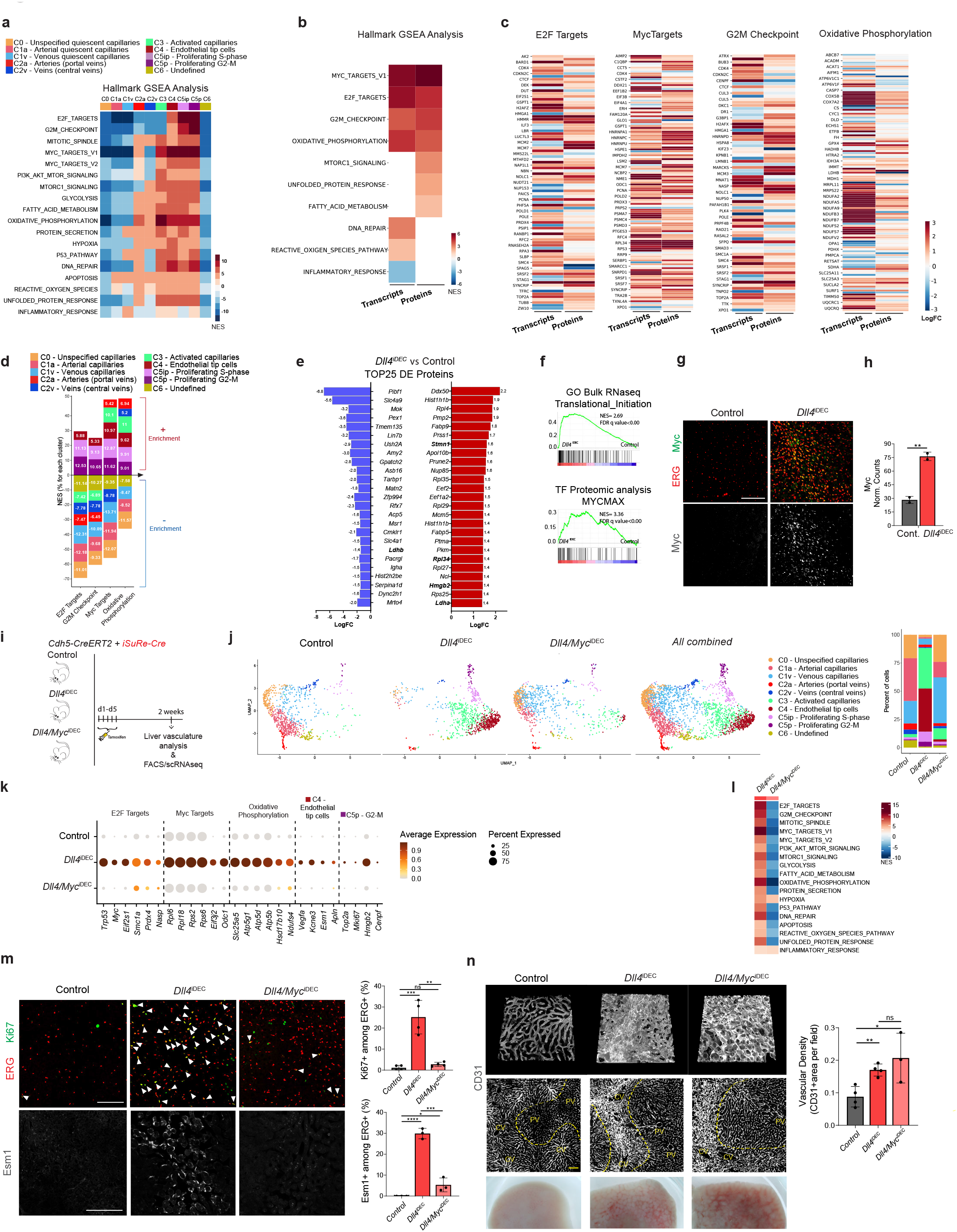
Myc loss prevents the Dll4^KO^ endothelial activation and single cell states but not vascular pathology. **a,** GSEA Hallmark analysis for each single cell cluster. **b,** GSEA Hallmark analysis performed with the *Dll4^iDEC^* bulk proteome and transcriptome. **c,** Heatmaps showing logFC of genes and proteins belonging to different sets. **d,** Barplot showing the Normalized Enrichment Score (NES) in each single cell cluster for the indicated gene sets. **e,** Barplot with the top differentially expressed (DE) proteins in *Dll4^iDEC^* livers. **f,** GSEA analysis show a significant positive enrichment in translational initiation-related genes and proteins encoded by genes that are regulated by the MycMax transcription factor binding site as shown by the Normalized Enrichment Score (NES). **g,** Micrographs with immunostainings showing Myc protein is upregulated in liver ECs (ERG+ cells) after *Dll4* deletion. **h,** *Myc* mRNA expression (normalized counts from bulkRNAseq). **i,** Experimental layout for the inducible deletion of *Dll4* and *Myc* in Cdh5+ ECs and scRNAseq analysis. **j,** UMAPs and barplot showing the 10 identified clusters and the percentage of cells for each cluster in the different samples. **k,** Dot plot of the top upregulated genes in *Dll4^iDEC^* liver ECs belonging to the indicated gene marker groups. **l,** GSEA Hallmark analysis showing the decreased expression of most gene sets in *Dll4/Myc^iDEC^*. **m,** Double deletion of *Dll4* and *Myc* in ECs results in a significant reversion of proliferation (Ki67+ERG+ cells) and Esm1+ expression (Esm1+ERG+) to control conditions. **n,** 3D Confocal micrographs from thick vibratome sections (top) or thin sections (bottom), and liver macroscopic pictures showing vessel enlargement and liver pathology in *Dll4/Myc^iDEC^* mutants similarly to *Dll4^DEC^* mutants. Charts n=2-4 animals per group minumum. Error bars indicate SD. *p < 0.05. **p < 0.01. ***p < 0.001. ****p < 0.0001. ns, non-significant.Scale bar, 100 μm.

We next investigated the implication of Myc in the *Dll4^iDEC^*transcriptional program and subsequent vascular-related pathology. Myc loss (in *Dll4/Myc^iDEC^* animals) almost entirely blocked the EC activation induced by *Dll4* loss, and very few cells were in the activated C3 and tip-cell C4 cluster (Fig.6i-k, Supplementary Fig. 5a-c). The exceptions were the hypoxia and inflammatory response pathways, which were still activated in *Dll4/Myc^iDEC^* ECs (Fig. 6l and Supplementary Fig. 5d). The frequencies of proliferating (Ki67+) and tip (Esm1+) cells in *Dll4/Myc^iDEC^*mutants were similar to those in wildtype animals (Fig. 6m). Myc activity is thus essential for the strong metabolic and biosynthetic phenotype of *Dll4^KO^* cells and the appearance of the abnormal cell states. Surprisingly, despite this strong transcriptional and cell states reversion to a quiescent state, *Dll4/Myc^iDEC^* mutant vessels were still highly abnormal and pathogenic (Fig. 6n). The vascular expansion in *Dll4/Myc^iDEC^* mutant livers was not in accordance with their more quiescent scRNAseq profile (Fig.6j-l), nor with the significantly lower frequencies of Ki67+ and Esm1+ cells (Fig. 6m). These data suggest that abnormal vascular expansion and pathology in *Dll4* mutant livers is not driven by the highly proliferative and tip-cell angiogenic states. Something else besides the switch in transcriptional identity and cell states is responsible for the vascular malformations and pathology observed after targeting Dll4.

### Anti-VEGF abolishes the *Dll4^KO^*-induced vascular pathology but not its transcriptional program

Among the few GSEA Hallmark pathways whose upregulation in *Dll4* mutants was not altered in *Dll4/Myc^iDEC^* vessels was the hypoxia pathway and inflammatory response (Fig. 6l). Hypoxia is known to induce expression of VEGFA, which can induce vascular expansion without the need for proliferation^38^. The expression of VEGFA was significantly upregulated in the *Dll4^KO^* tip cell cluster (Sup. Fig. 6a). Therefore, we explored if anti-VEGF would be able to prevent the appearance of the activated vascular cell states and liver pathology induced by *Dll4* deletion. Unlike Myc loss, anti-VEGF treatment reduced both the vascular expansion and the liver pathology induced by *Dll4* deletion (Fig. 7a-d, Supplementary Fig. 6b). scRNAseq analysis confirmed the almost complete loss of the tip-cell (C4) and proliferating (C5) single-cell states, as well as a significant reduction in the activated cell states (C3), with a general return to the quiescent cell states (Fig. 7e-g, Supplementary Fig. 6c). The loss of the arterial capillary cluster (C1a) after Dll4 targeting was, however, not prevented by anti-VEGF (Fig. 7e, 7g and Supplementary Fig. 6d, e). The scRNAseq data also revealed an increase in the relative number of ECs from larger veins (central veins) and arteries (portal veins), suggesting a depletion of VEGFR2/Kdr+ sinusoidal capillaries by anti-VEGFA (Fig. 7e and Supplementary Fig. 6a Vegfr2/Kdr expression). Liver sinusoidal capillaries density and EC numbers were also significantly lower than at baseline in wildtype livers (Fig. 7b,d,i). Anti-VEGFA also prevented the upregulation of *Myc* and its target *Odc1* and several other cell cycle or tip-cell related genes observed in *Dll4^iDEC^* mutant ECs (Fig. 7g-i and Supplementary Fig. 6f).

**Figure 7:**
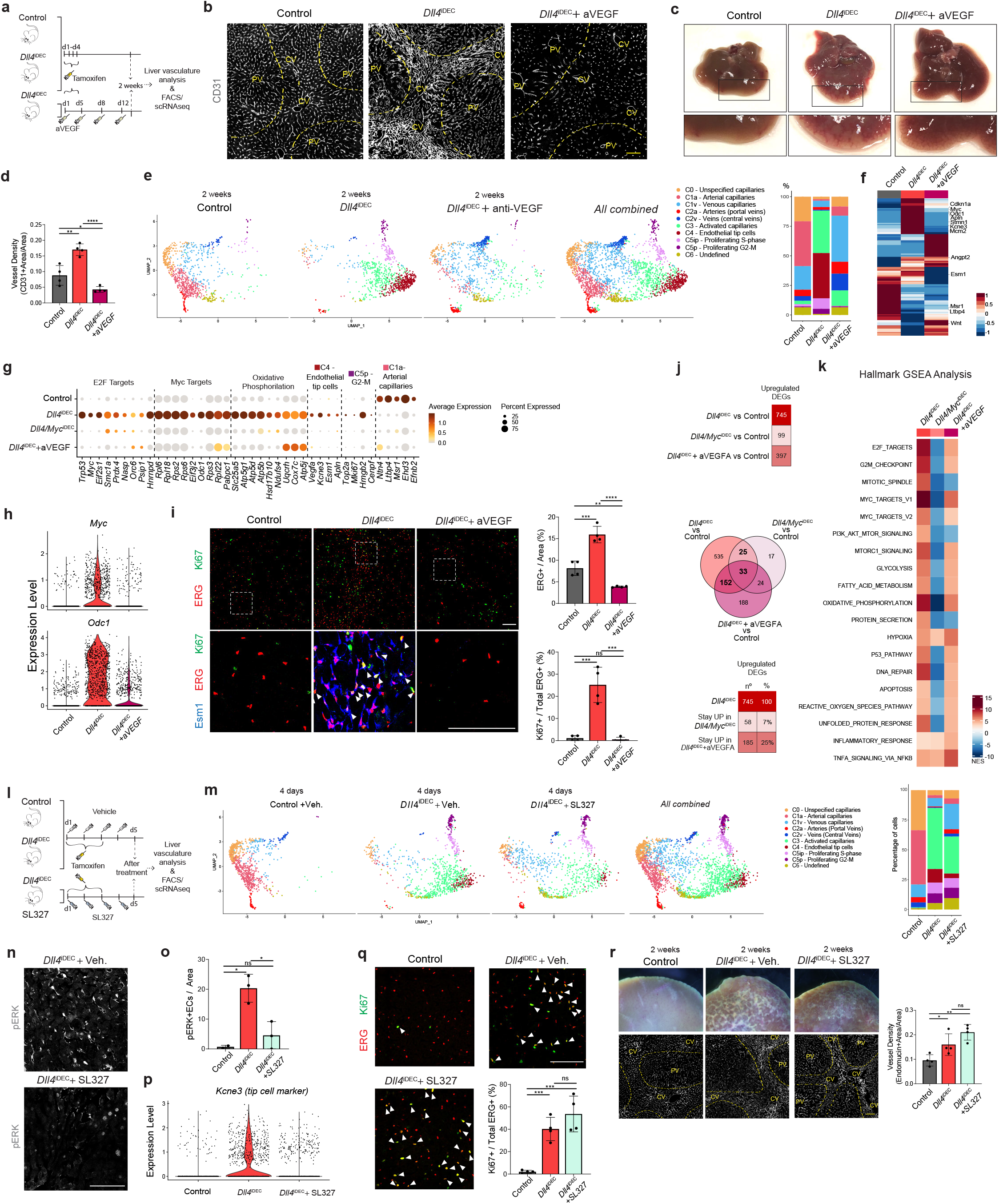
Vascular abnormalities and liver pathology are prevented by anti-VEGFA administration in *Dll4^iDEC^* mutants by ERK-independent mechanisms. **a,** Experimental layout for the inducible deletion of *Dll4* in Cdh5+ ECs and anti-VEGFA administration. **b,** Confocal micrographs showing reduced CD31/vascular immunostaining. **c,** Macroscopic liver pictures of indicated samples. **d,** Vessel density (CD31) is reduced in *Dll4^iDEC^* mutants after anti-VEGFA. **e,** UMAPs and barplot showing the 10 identified clusters and the percentage of cells for each cluster in indicated samples. **f,** Unsupervised hierarchical clustering showing gene expression changes. **g,** Dot plot of the top upregulated genes for each indicated gene set. **h,** Violin plots of scRNAseq data showing that Anti-VEGF prevents the strong upregulation of *Myc* and its target *Odc1*. **i,** The total number of ERG+ ECs, proliferation (Ki67+ERG+) and Esm1 expression (Esm1+ERG+) is rescued to control conditions after anti-VEGF administration. **j,** Number of upregulated genes for each contrast and Venn diagrams showing that when compared with Myc loss, Anti-VEGF has a lower impact on the *Dll4^iDEC^* upregulated genetic programme. **k,** GSEA Hallmark analysis confirms the more moderate effect of Anti-VEGF on the *Dll4^iDEC^* genetic programme when compared with Myc loss. **l,** Experimental layout for the inducible deletion of Dll4 and SL327 administration. **m,** UMAPs and barplot showing the 10 identified clusters and the percentage of cells for each cluster in indicated samples. **n-o,** The administration of an ERK/MEK signalling inhibitor (SL327) results in lack of ERK phosphorylation. **p,** Violin plot showing that SL327 treatment inhibits the generation of tip cells (Kcne3+). **q,** The administration of SL327 does not change the frequency of proliferating Ki67+ ECs (Ki67+ERG+).**r,** Abnormal vasculature (CD31+) associated to liver pathology still occurs after SL327. Error bars indicate SD. *p < 0.05. **p < 0.01. ***p < 0.001. ****p < 0.0001. ns, non-significant. Scale bars, 100 μm.

These results show that anti-VEGF prevents not only the appearance of the abnormal single-cell states induced by Dll4 targeting, as Myc loss also does, but in addition it also prevents the vascular expansion and associated organ pathology. However, blocking VEGF had a much lesser impact than Myc loss on the *Dll4^KO^*-upregulated gene signature (Fig. 7j). Anti-VEGFA treatment of *Dll4^iDEC^* livers attenuated, but did not completely downregulate, many of the genes associated with Myc and its biosynthetic and proliferation activities, including oxidative phosphorylation and RNA processing (Fig. 7k, Supplementary Fig. 6g, h). This suggests that even though *Dll4^KO^* + Anti-VEGF-treated ECs are more metabolically active than *Dll4/Myc^iDEC^* ECs, they do not result in liver vascular abnormalities and pathology.

VEGFA induces many important and often difficult to distinguish endothelial functions, such as proliferation, sprouting, survival, and permeability^39–41^. VEGF is thought to execute its effects on sprouting and angiogenesis mainly through ERK signaling^42,43^. However, administration of a highly effective ERK/MEK signaling inhibitor (SL327) had a much more modest effect than anti-VEGF (Fig. 7l-o). SL327 inhibited mainly the generation of tip cells (*Kcne3+* and cluster C4 in Fig.7m and 7p) and did not change the frequency of proliferating (Ki67+) ECs (Fig. 7q), and organ pathology still occurred (Fig. 7r top). Interestingly, SL327 also did not prevent the vascular expansion of *Dll4^iDEC^* liver and retina vessels (Fig. 7r bottom and Supplementary Fig. 6i). Thus, the vascular pathophysiology effects of anti-VEGF, and also anti-Dll4, are broader than and independent of ERK signaling. This is in line with the absence of major vascular abnormalities in *Notch1^iDEC^ and Rbpj^iDEC^* mutant vessels, despite the higher ERK signaling than in *Dll4^iDEC^* mutants.

Overall, these results show that the genetic and pharmacological modulation of single-cell states related with endothelial dedifferentiation, activation, proliferation and sprouting often do not correlate with vascular phenotypes and ultimately organ pathology. The transcriptional changes and angiogenic cell states elicited by targeting Dll4 are not the cause of its vascular pathophysiology. Instead, we propose that is the unrelated vascular enlargement and malfunction that leads to organ pathology. Therefore, vascular cell states arising from single cell transcriptomics should not be used as markers of vascular expansion, function and ultimately organ pathophysiology.

## Discussion

Notch is one of the most important pathways for vascular development by enabling the necessary differentiation of endothelial cells during angiogenesis^44,45^. Here, we show that Notch also plays an important role in the homeostasis of several organ vascular beds. Dll4 is active in all organ vascular beds, and its loss impacts the transcriptome of most quiescent ECs; however, Dll4 targeting only effectively activates vascular growth in the heart, muscle, and liver. Even though the existence of 5 ligands and 4 receptors in the Notch pathway allows for the possibility of multiple quantitative and qualitative signaling combinations and redundancy, our results confirm that Dll4 and Notch1 are clearly the most important Notch ligand-receptor pair for maintaining the global homeostasis of ECs.

Previous work suggested that Dll4 and Notch1/Rbpj have similar functions in vascular development and homeostasis^12,13,35,36, 46–51^, with only Jagged ligands shown to have opposite functions in Notch signaling and angiogenesis^36^. This is the first study showing that Dll4 can have distinct functions from its receptors in vascular biology. It was possible to identify this difference only thanks to the use of scRNAseq and high-resolution confocal analysis of vessels morphology; bulkRNAseq analysis did not reveal significant differences between the transcriptomes of Dll4 and Rbpj mutants. Dll4 targeting, unlike Notch receptor or Rbpj targeting, elicits a unique cascade of changes that culminates in the loss of not only the arterial capillary cell state, but also of the quiescent venous capillary cell state, with a significant increase in the proportion of highly activated cells with either tip-cell or proliferating cell signatures. Paradoxically, although Dll4 loss induces a weaker loss of Notch signaling than is induced by the loss of Notch receptors or Rbpj, it elicits a much stronger metabolic activation and expansion of the endothelium. This may be in part related to the bell-shaped response of ECs to mitogenic stimuli, as we previously showed during retina angiogenesis^29^. Our data indicate that full loss of Notch, or Rbpj, induces stronger ERK signaling and hypermitogenic arrest associated with hallmarks of cellular senescence, whereas *Dll4^iDEC^*vessels retain a residual level of Notch signaling, that instead effectively induces strong Myc-driven ribosome biogenesis and a metabolic switch towards active protein synthesis and cell growth that drives both EC proliferation and the generation of tip cells. The pro-proliferative and pro-sprouting effect of targeting Dll4 in quiescent vessels is in contrast to the hypermitogenic cell-cycle arrest that occurs after targeting Dll4 during embryonic and retina angiogenesis^29,52^, presumably a reflection of the significantly lower levels of growth factors, including VEGF, in adult organs.

High resolution confocal microscopy revealed the heterozonal effect of Dll4 targeting. The induction of EC proliferation and tip cells was restricted to the most hypoxic liver venous sinusoids, precisely the ones with lower expression of Dll4 and Notch. Previous research showed that liver venous sinusoids have higher baseline activity of several tyrosine kinase signaling pathways^53^, which may explain the observed zonal effect of Dll4 targeting.

The temporal analysis of the effects of Dll4 targeting on the adult liver vasculature also revealed that it takes at least a week for the full transcriptional reprogramming of quiescent ECs and the vascular expansion and organ pathology to become noticeable. During angiogenesis, this transcriptional and vascular morphology switch is already evident after 24 hours of anti-Dll4 treatment^29^. This slow transcriptional reprogramming of quiescent ECs by Dll4 targeting may be related with the much lower levels of growth factors and nutrients availability in adult organs. The slow nature of this reprogramming may also explain the lack of effect of the small molecule inhibitor DBZ on quiescent ECs. Unlike anti-Dll4 or genetic deletion, which result in continuous loss of signaling, the less stable small-molecule inhibitor DBZ elicited no significant change in the quiescent vascular cell transcriptional states and phenotypes, whereas it is very effective during retina angiogenesis^29,46^.

These findings have implications for selecting the most effective and safe way to target Notch in the clinic, with blocking antibodies targeting Dll4 versus antibodies targeting Notch receptors, or the use of small-molecule inhibitors. Our data suggest that Notch-receptor targeting antibodies or small-molecule γ-secretase inhibitors will be as effective as anti-Dll4 at dysregulating tumour or ischemia-related angiogenesis, but far less effective at changing quiescent vascular single-cell states and less likely to induce vascular pathology, which can be beneficial in some therapeutic settings.

Our analysis also confirms the importance of Myc for the biology of ECs in the absence of Dll4. We previously reported that *Myc* loss rescues the ability of *Rbpj^KO^* or *Dll4^KO^* ECs to form arteries^52^. Here, we show that Myc loss abrogates the generation of activated, proliferative, and sprouting tip cells after Dll4 targeting, but surprisingly, this return to genetic and phenotypic quiescence is insufficient to prevent Dll4-targeting–induced vascular expansion, dysfunction, and consequent organ pathology. In contrast, anti-VEGF treatment did not completely abrogate the Dll4-targeting genetic program, but was able to prevent most *Dll4^iDEC^* single cell states and the associated vascular expansion, dysfunction and organ pathology. This effect of anti-VEGF was independent from its canonical function on ERK signaling and cell-sprouting, as it was not reproduced by the use of a MEK/ERK inhibitor. These data point towards a broader role for VEGF in vascular expansion and survival, that is independent of its effect on cell proliferation and sprouting, as previously proposed^38,54^.

Altogether, our data obtained with several compound mutant and pharmacological approaches (Supplementary Fig.7) show that the vascular dysfunction and organ pathology that follows Dll4 targeting is induced not by the loss of arterial cell states or the gain of proliferative and tip-cell states, but rather by the transcriptionally unrelated or undetectable expansion and malfunction of the vascular network. These data raises questions about the general use of single cell transcriptional or genetic states to describe and predict functional or dysfunctional vascular phenotypes and ultimately organ pathology. It also sheds light on the best strategies to therapeutically modulate angiogenesis, reprogram endothelial quiescent cell states, or avoid vascular-related organ pathology with pharmacological compounds targeting these pathways.

## MATERIALS AND METHODS

### Mice

The following mouse lines were used and interbred: *Tg(Cdh5-CreERT2)*^55^, *Tg(iSuRe-Cre)*^32^, *Dll1^flox/flox^* ^56^, *Jag1^flox/flox^* ^57^, *Jag2^flox/flox^* ^58^, *Dll4^flox/flox^* ^59^, *Notch1^flox/flox^* ^60^, *Notch2^flox/flox^*^61^, *Notch4^KO^* (generated as described below), *Rbpj^flox/flox^*^62^, *Myc^flox/flox^* ^63^, *Cdkn1a(p21)^KO^* ^64^, *iChr-Mosaic* ^65^ and *iMb-Mosaic* ^65^. To induce CreERT2 activity in adult mice, 20 mg or 10 mg tamoxifen (Sigma-Aldrich, T5648) were first dissolved in 140 µl absolute ethanol and then in 860 µl corn oil (20 mg/ml or 10 mg/ml tamoxifen). This solution was given by intraperitoneal injection (200 µl, total dose of 1 or 2 mg tamoxifen, respectively) to adult mice (8-12 weeks old). For *Dll4*, *ifgMosaic,* or short-term inducible loss-of-function experiments, 1 mg tamoxifen was injected on 3 consecutive days. For all other inducible loss-of-function experiments, 2 mg tamoxifen was injected for 5 consecutive days. To activate recombination in pups containing CreERT2 alleles, tamoxifen was injected at the indicated stages at a dose of 40 ug/g body weight. All mouse lines and primer sequences required to genotype these mice are provided in Table 1.

**Table 1:**
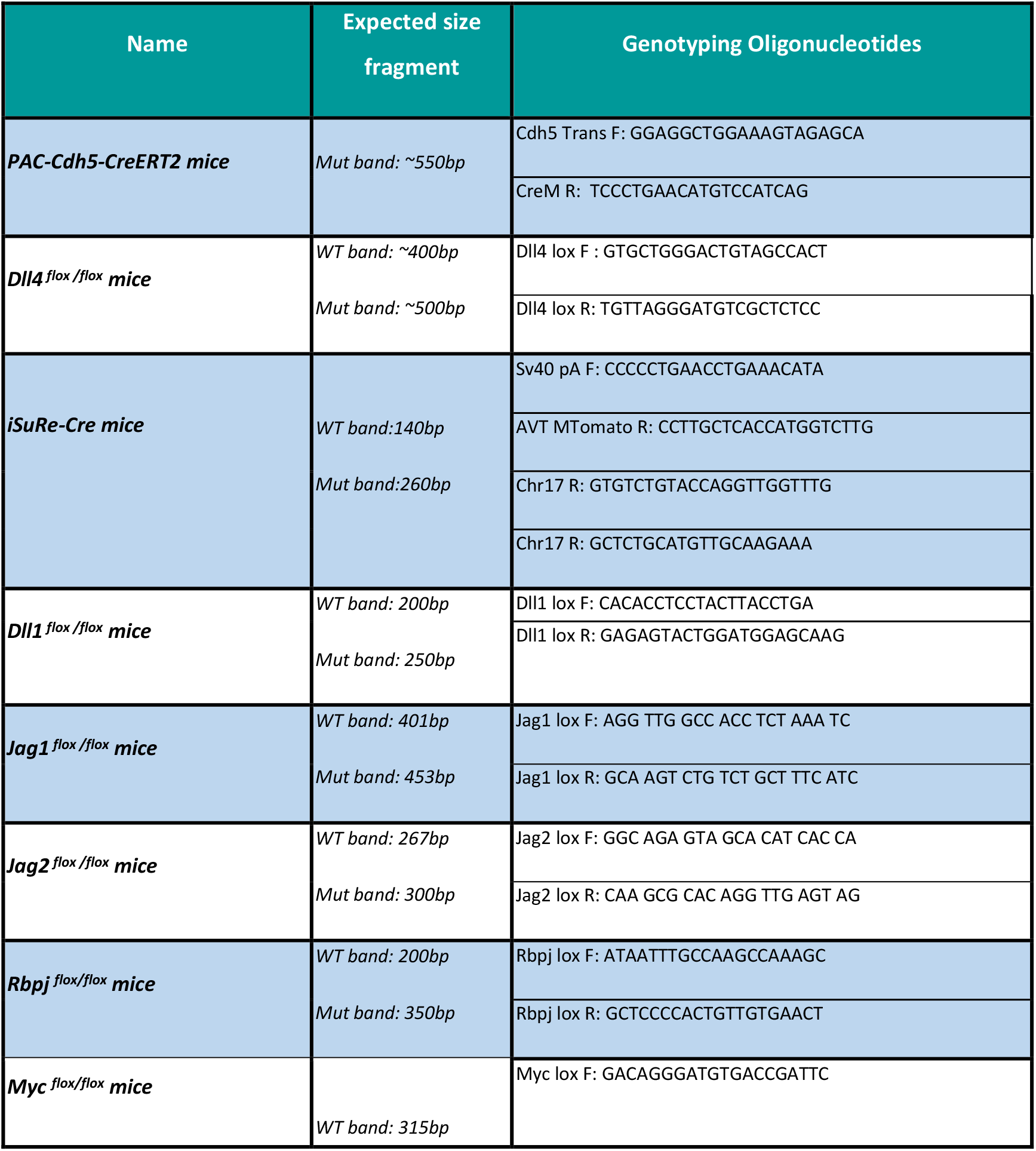
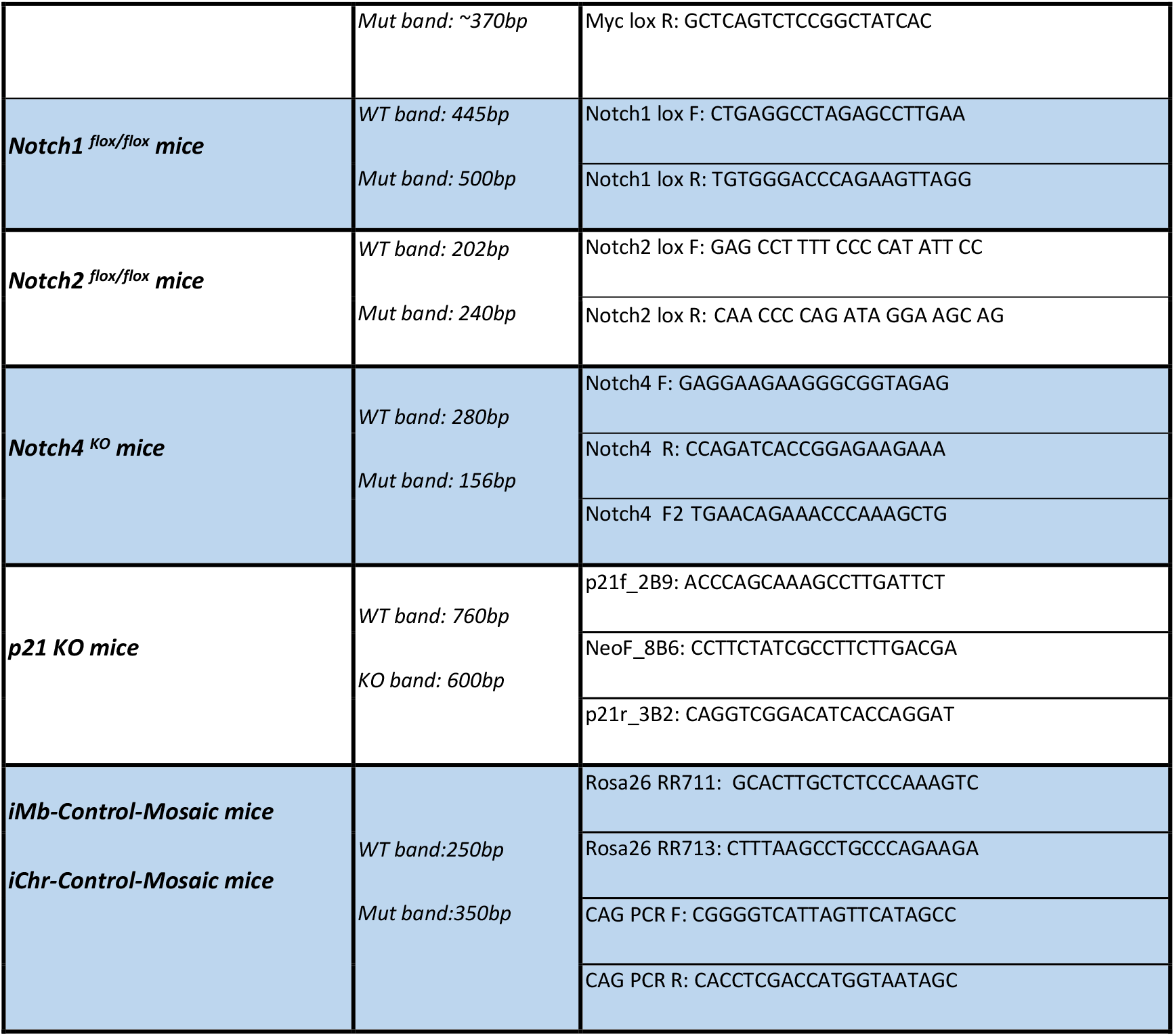
Mouse lines and genotyping PCR primers.

Dll4/Notch signaling inhibition in ECs was achieved using blocking antibodies to murine Dll4 (REGN1035), developed by Regeneron. Mouse IgG (Sigma) was used in littermates as a control treatment. Mice received a single intraperitoneal injection of 200 µl IgG or anti-Dll4 (20 mg/kg in PBS). Livers were collected 48h after blocking antibody administration and processed for immunohistochemistry and bulk-RNAseq. The γ-secretase inhibitor DBZ (YO-01027, S2711, Selleckchem) was injected intraperitoneally at 30µmol/kg on 5 consecutive days. The last injection was administered 16h before mice were sacrificed and livers collected for scRNAseq analysis. To inhibit ERK phosphorylation, 150µl SL327 (MEK inhibitor from Selleckchem, at 120 mg/kg) was injected on 5 or 14 consecutive days. The final injections were administered 2 h before sacrifice. For anti-VEGFA experiments, mouse Anti-VEGFA G6-31, developed by Genentech, was administered four times over 14 days at a concentration of 5 mg/kg.

All mouse husbandry and experimentation was conducted using protocols approved by local animal ethics committees and authorities (Comunidad Autónoma de Madrid and Universidad Autónoma de Madrid CAM-PROEX 177/14 and CAM-PROEX 167/17). The CNIC mouse colony (Mus musculus) is maintained in racks with individual ventilation cages according to current Spanish and European legislation (RD 53/2013 and EU Directive 63/2010). Mice have dust/pathogen-free bedding, and sufficient nesting and environmental enrichment material for the development of species-specific behavior. All mice have ‘ad libitum’ access to food and water in environmental conditions of 45–65% relative humidity, temperatures of 21–24 °C, and a 12 h/12 h light/dark cycle. In addition, and to preserve animal welfare, mouse health is monitored with an animal health surveillance program, which follows FELASA recommendations for specific pathogen-free facilities.

We used Mus musculus with C57BL6 or C57BL6×129SV genetic backgrounds. To generate mice for analysis, we intercrossed mice with an age range between 7 and 30 weeks. All adult mice analyzed were less than 12 weeks old. We do not expect our data to be influenced by mouse sex.

To generate Notch4^KO^ mice we used guide RNAs Notch4_1 (agggaccctcagagcccttg) and Notch4_2 (agggaatgatgccacgcata) to target mouse Notch4 in mouse eggs from the C57Bl6 genetic background. Injection mixture was composed by the described crRNAs (IDT) and tracrRNA (IDT, catalog 1072533) at 0.305 microM and Cas9 nuclease (Alt-R® S.p. HiFi Cas9 Nuclease V3, 100 µg, Catalog 1081060) at 20 ng/microL.

Founders were screened by PCR with the primers below to confirm the genetic deletion.

### Immunofluorescence on cryosections

For immunostaining on organ cryosections, tissues were fixed for 2 h in 4% PFA in PBS at 4°C. After three washes in PBS for 10 min each, organs were stored overnight in 30% sucrose (Sigma) in PBS. Organs were then embedded in OCT^TM^ (Sakura) and frozen at −80°C. Cryosections (35μm) were cut on a cryostat (Leica), washed three times for 10 min each in PBS, and blocked and permeabilized in PBS containing 10% donkey serum (Milipore), 10% fetal bovine serum (FBS), and 1% Triton X-100. Primary antibodies were diluted in blocking/permeabilization buffer and incubated overnight at 4°C. This step was followed by three 10 min washes in PBS and incubation for 2 h with conjugated secondary antibodies (1:200, Jackson Laboratories) and DAPI in PBS at room temperature. After three washes in PBS, sections were mounted with Fluoromount-G (SouthernBiotech). All antibodies used are listed in Table 2. To detect Ki67 or c-Myc in the same section as ERG, rabbit anti-Ki67 or anti-cMyc was used together with a Fab fragment CY3 secondary antibody, which is compatible with the later use of rabbit anti-ERG-Alexa647.

**Table 2:** Antibodies list.

| ANTIBODIES | DILUTION | SOURCE | IDENTIFIER |
| --- | --- | --- | --- |
| Anti-GFP/YFP/Cerulean | 1 to 200 (IF) | Acris Antibodies | Cat# R1091P |
| Anti-DsRed | 1 to 400 (IF) | Clontech | Cat# 632496 |
| Anti-HA -647 | 1 to 200 (IF) | Cell Signaling Technology | Cat# 3444S |
| Anti-ERG | 1 to 400 (IF) | Abcam | Cat# ab110639 |
| Anti-ERG-AF-647 | 1 to 200 (IF) | Abcam | Cat# ab196149 |
| Anti-Ki67 | 1 to 200 (IF) | Thermo Fisher | Cat# RM-9106-S0 |
| Anti-Endomucin | 1 to 200 (IF) | Santa Cruz Biotechnology | Cat# SC-53941 |
| Anti-CD31 | 1 to 200 (IF) | BD Biosciences | Cat# 553370 |
| Anti-p21 | 1 to 100 (IF) | Santa Cruz Biotechnology | Cat# SC-397-G |
| Anti-Phospho-ERK | 1 to 100 (IF)<br>1 to 1000 (WB) | Cell Signaling Technology | Cat# 4370S |
| Anti-Myc | 1 to 200 (IF) | Millipore | Cat# 06-340 |
| Anti-cleaved N1ICD | 1 to 200 (IF) | Cell Signaling Technology | Cat# 4147 |
| Anti-Dll4 | 1 to 200 (IF) | R&D system | Cat# AF1389 |
| Anti-alpha/beta-Tubulin | 1 to 5000 (WB) | Cell Signaling Technology | Cat# 2148 |
| Anti-PhiYFP | 1 to 200 (IF) | Axxora | Cat# EVN-AB605 |
| Anti-Jagged1 | 1 to 100 (IF) | Cell Signaling Technology | Cat# 2620 |
| Anti-VEGFR2 | 1 to 200 (IF) | BD Pharmingen | Cat# 550549 |
| Anti-CD34 | 1 to 200 (IF) | BD Biosciences | Cat# 560238 |
| Anti-CD45 | 1 to 200 (IF) | TonboBio | Cat# 35-0454-U100 |
| Anti-Caspase 3 | 1 to 50 (IF) | Cell Signaling Technology | Cat# 9661S |
| Anti-Esm1 | 1 to 200 (IF) | R&D system | Cat# AF1999 |
| Anti-Msr1 | 1 to 200 (IF) | R&D system | Cat# AF1797-SP |
| Alexa fluor-conjugated antibodies | 1 to 400 (IF) | Life Technologies | N/A |

### Vibratome immunofluorescence and 3D imaging

For immunostaining on vibratome sections, tissues were fixed in 4% PFA (in PBS) and washed as above. Organs were then embedded in 6% agarose low melting gel (Invitrogen), and organ sections (100 μm) were cut on a vibratome. Sections were permeabilized for 1 h in PBS containing 1% Triton X-100 and 0.5% Tween20. Sections were then blocked for 1 h in a PBS solution containing 1% Triton X-100, 10% donkey serum, and 10% FBS. Primary antibodies were diluted in blocking buffer and incubated with sections overnight at 4°C. This step was followed by 6 washes with 1% Triton X-100 in PBS for 15 min and incubation for 2 h with conjugated secondary antibodies (1:200, Jackson Laboratories) and DAPI in PBS at room temperature. After three 15 min washes in PBS, sections were mounted in Fluoromount-G (SouthernBiotech). All antibodies used are listed in Table 2.

### Immunofluorescence on paraffin sections

The short-lived and difficult to detect cleaved/activated N1ICD epitope and the Jag1 ligand were detected with the TSA amplification kit (NEL774) procedure in paraffin sections after antigen retrieval. Briefly, sections were dewaxed and rehydrated, followed by antigen retrieval in sub-boiling sodium citrate buffer (10mM, PH=6.0) for 30 min. The slides were cooled down to RT for 30min, followed by incubation for 30 minutes in 3% H_2_O_2_ in methanol to quench endogenous peroxidase activity. Next, slides were rinsed in double-distilled H_2_O and washed for three times 5 min in PBS, followed by blocking for 1 h in PBS containing 3% BSA, 200mM MgCl_2_, 0.3% Tween20, and 5% donkey serum. Sections were then incubated with primary antibody in the same solution overnight at 4°C. After washes, slides were incubated for 2 h with anti-rabbit-HRP secondary antibody at RT, and, after washing, the signal was amplified using the TSA fluorescein kit (NEL774). Sections were mounted with Fluoromount-G (SouthernBiotech). All antibodies used are listed in Table 2.

### In vivo EdU labelling and EC proliferation detection

To detect EC proliferation in adult livers, 20 μg g^−1^ body weight EdU (Invitrogen, A10044) was injected intraperitoneally into adult mice 5 h before dissection. Livers were isolated for cryosection analysis. EdU signals were detected with the Click-it EdU Alexa Fluor 647 or 488 Imaging Kit (Invitrogen, C10340 or C10337). In brief, after all other primary and secondary antibody incubations, samples were washed according to the immunofluorescence staining procedure and then incubated with Click-iT EdU reaction cocktail for 40 min, followed by DAPI counterstaining.

### Image acquisition and analysis

Immunostained organ sections were imaged at high resolution with a Leica SP5, SP8, or SP8 Navigator confocal microscopes fitted with 10x, 20x, or 40x objectives for confocal scanning. Individual fields or tiles of large areas were acquired from cryosections and paraffin sections. Large Z-volumes of the vibratome samples were imaged for 3D representations. All images shown are representative of the results obtained for each group and experiment. Animals were dissected and processed under exactly the same conditions. Comparisons of phenotypes or signal intensity were made with pictures obtained using the same laser excitation and confocal scanner detection settings. Fiji/ImageJ was used to threshold, select, and quantify objects in confocal micrographs. Both manual and automatic ImageJ public plugins and customed Fiji macros were used for quantification.

### Western blot analysis

For the analysis of protein expression, livers were transferred to a reagent tube and frozen in liquid nitrogen. On the day of the immunoblotting, the tissue was lysed with lysis buffer [Tris-HCl pH=8 20mM, EDTA 1mM, DTT 1mM, Triton X-100 1% and NaCl 150mM, containing protease inhibitors (P-8340 Sigma) and phosphatase inhibitors (Calbiochem 524629) and orthovanadate-Na 1 mM] and homogenized with a cylindrical glass pestle. Tissue and cell debris was removed by centrifugation, and the supernatant was diluted in loading buffer and analyzed by SDS–PAGE and immunoblotting. Membranes were blocked with BSA and incubated with primary antibodies listed in Table 2.

### Endothelial cell isolation for transcriptomic and proteomic analysis

The following methods were used to isolate ECs for qRT-PCR, bulk-RNAseq, and proteomics analysis. To profile quiescent ECs with full loss of Dll4, tamoxifen was injected on 3 consecutive days into *Tg(Cdh5-CreERT2)*; *Dll4^flox/flox^* mice. To analyze loss of *Notch1* and *Rbpj*, tamoxifen was injected on 5 consecutive days into *Tg(Cdh5-CreERT2) Tg(iSuRe-Cre*); *Notch1^flox/flox^* mice and *Tg(Cdh5-CreERT2); Tg(iSuRe-Cre); Rbpj^flox/flox^* ^62^ mice. Controls were tamoxifen-injected C57BL/6J mice and *Tg(Cdh5-CreERT2); Tg(iSuRe-Cre*) mice respectively. At day 14 after the first tamoxifen injection, heart, lungs, liver, and brain were dissected, minced and digested with 2.5 mg/ml collagenase type I (Thermofisher), 2.5 mg/ml dispase II (Thermofisher), and 50 ng/ml DNAseI (Roche) at 37°C for 30 min to create a single-cell suspension. Cells were passed through a 70 µm filter to remove non-dissociated tissue. Erythroid cells were removed from cell suspensions by incubation with blood lysis buffer (0.15 M NH4Cl, 0.01M KHCO3, and 0.01 M EDTA in distilled water) for 10 min on ice. Cell suspensions were blocked in blocking buffer (DPBS containing no Ca2+ or Mg2+ and supplemented with 3% dialyzed FBS; Thermo Fisher). For EC analysis, cells were incubated at 4°C for 30 min with APC-conjugated rat anti-mouse CD31 (1:200, BD Pharmigen, 551262). DAPI (5 mg/ml) was added to the cells immediately before FACS, which was performed with FACS Aria (BD Biosciences) or Synergy4L cell sorters. For the isolation of ECs from dissociated tissues, viable cells were selected by DAPI-negative fluorescence. All viable cells were interrogated by examining FSC and SSC to select for size and complexity, and by comparing FSC-H and FSC-W repeated with SSC-H and SSC-W to discern single cells. An additional channel lacking any fluorescent label was also acquired to detect and exclude autofluorescence. Cells were selected by their APC and/or endogenous MbTomato positive signal and were gated with a negative control sample. For qRT-PCR and bulk-RNAseq experiments, approximately 10000-20000 cells for each group of DAPI-negative APC-CD31+ ECs (for Dll4 loss of function and control), DAPI negative APC-CD31+/MbTomato+ ECs (for Notch1 or Rbpj loss of function and control) were sorted directly to RLT buffer (RNAeasy Micro kit - Qiagen). RNA was extracted with the RNAeasy Micro kit and stored at −80°C. For proteomic analysis, approximately 3×10^6^ DAPI-negative APC-CD31+ ECs per group were sorted directly to blocking buffer. Cells were spun down for 10 min at 350g and pellet stored at −80°C.

To isolate ECs for scRNAseq experiments, 1.5 mg tamoxifen was injected on 4 consecutive days into Control (*Tg(Cdh5-CreERT2)*, *Tg(iSuRe-Cre)*); *Dll4^iDEC^* (*Dll4^flox/flox^*, *Tg(Cdh5-CreERT2)*, *Tg(iSuRe-Cre); Notch1^iDEC^* (*Notch1^flox/flox^, Tg(Cdh5-CreERT2, Tg(iSuRe-Cre)*; *Rbpj^iDEC^* (*Rbpj^flox/flox^, Tg(Cdh5-CreERT2, Tg(iSuRe-Cre); Dll4^Het^ (Dll4^flox/wt^*, *Tg(Cdh5-CreERT2)*; *Tg(iSuRe-Cre)); Notch1/2/4^iDEC^* (*Notch1/2/4^flox/flox^*, *Tg(Cdh5-CreERT2)*; *Tg(iSuRe-Cre)); Jag1/Jag2/Dll1^iDEC^ (Jag1/2/Dll1^flox/flox^*, *Tg(Cdh5-CreERT2)*; *Tg(iSuRe-Cre); Dll4/Myc^iDEC^* (*Dll4/Myc^flox/flox^*, *Tg(Cdh5-CreERT2)*, *Tg(iSuRe-Cre))*. At day 14 after the first tamoxifen injection, livers were dissected, minced, and digested for 30 min with pre-warmed (37C) dissociation buffer (2.5mg/ml collagenase I (Thermo Fisher 17100017), 2.5mg/ml dispase II (Thermo Fisher 17105041), 1ul/ml DNAse in PBS containing Ca2+ and Mg2+ (Gibco)). The digestion tube was agitated every 3-5 minutes in a water bath. At the end of the 30 minutes incubation, sample tubes were filled up to 15 ml with sorting buffer (PBS containing no Ca2+ or Mg2+ and supplemented with 10% FBS (Sigma, F7524)) and centrifuged (450g, 5 min, 4°C). The supernatant was aspirated, and cell pellets were resuspended in 2ml 1x RBC lysis buffer (BioLegend, 420301) and incubated for 5 min on ice. To each sample were added 6 ml of sorting buffer, and samples were then passed through a 70um filter. Live cells were counted in a Neubauer Chamber using trypan blue exclusion. Cells from each condition (4×10^6^/condition) were collected in separate tubes, and cells were incubated for 30 min with horizontal rotation in 300ul antibody incubation buffer (PBS + 1% BSA) containing 1 µl CD31-APC, 1 µl CD45-APC-Cy7, and 1µl of hash tag oligo (HTO) conjugated antibodies (Biolegend). HTOs were used to label and distinguish the different samples when loaded on the same 10x genomics port, and in this way also guarantee the absence of batch effects. After the antibodies incubation, samples were transferred to 15 ml Falcon tubes, 10 ml sorting buffer was added, and samples were centrifuged (450g, 5min, 4°C). The supernatant was aspirated, pellets were resuspended in 1.5 ml sorting buffer and transferred to Eppendorf tubes, and the resulting suspensions were centrifuged (450g, 5min, 4°C). The resulting pellets were resuspended in 300 µl sorting buffer containing DAPI. Cells were sorted by FACS with an Aria Cell Sorter (BD Biosciences); ECs were identified as described below, and CD31+, CD45-MbTomato+ cells were sorted.

### Next Generation Sequencing sample and library preparation

Next Generation Sequencing (NGS) experiments were performed in the Genomics Unit at CNIC. For bulk-RNAseq experiments the pipeline performed was the following. For *Dll4*^iDEC^ ECs samples, 1 ng of total RNA was used to amplify the cDNA using the SMART-Seq v4 Ultra Low Input RNA Kit (Clontech-Takara) following manufactureŕs instructions. Then, 1 ng of amplified cDNA was used to generate barcoded libraries using the Nextera XT DNA library preparation kit (Illumina). Basically, cDNA is fragmented and adapters are added in a single reaction followed by an amplification and clean up. The size of the libraries was checked using the Agilent 2100 Bioanalyzer High Sensitivity DNA chip and their concentration was determined using the Qubit® fluorometer (ThermoFisher Scientific). Libraries were sequenced on a HiSeq2500 (Illumina) to generate 50 bases single end reads. FastQ files for each sample were obtained using CASAVA v1.8 software (Illumina). For *Rbpj*^iDEC^ and *Notch1*^iDEC^ ECs samples, between 400 and 3000 pg of total RNA were used to generate barcoded RNA-seq libraries using the NEBNext Single Cell/Low Input RNA Library Prep Kit for Illumina (New England Biolabs) according to manufacturer’s instructions. First cDNA strand synthesis was performed, then cDNA was amplified by PCR followed by fragmentation. Next, cDNA ends were repaired and adenylated. The NEBNext adaptor was then ligated followed by second strand removal, uracile excision from the adaptor and PCR amplification. The size of the libraries was checked using the Agilent 2100 Bioanalyzer and the concentration was determined using the Qubit® fluorometer (Life Technologies). Libraries were sequenced on a HiSeq2500 (Illumina) to generate 60 bases single reads. FastQ files for each sample were obtained using bcl2fastq 2.20 Software software (Illumina).

For scRNAseq experiments the pipeline performed was the following. Single cell suspensions of freshly isolated ECs were resuspended in Cell Capture buffer to a final concentration of 800 to 1600 cells/µl. scRNA-seq libraries were prepared using the Chromium Single Cell 3′ Reagent Kits v3.1 (10x Genomics; Pleasanton, CA, USA) according to the manufacturer’s instructions. The aimed target cell recovery for each port was 10000 cells, with a 2000-2500 aimed cell recovery per experimental condition labelled with a given hashtag antibody. Generated libraries were sequenced on an Illumina HiSeq4000 or NextSeq2000, followed by de-multiplexing.

### Transcriptomic data analysis

Transcriptomic data were analyzed by the CNIC Bioinformatics Unit.

For bulk RNA-seq, the number of reads per sample was between 12 and 42 million. Reads were processed with a pipeline that assessed read quality using FastQC (Babraham Insitute-http://www.bioinformatics.babraham.ac.uk/projects/fastqc/) and trimmed sequencing reads using cutadapt ^66^, eliminating Illumina and SMARTer adaptor remains and discarding reads < 30 bp. More than 93% of reads were kept for all samples. Resulting reads were mapped against the mouse transcriptomes GRCm38.76 and GRCm38.91, and gene expression levels were estimated with RSEM ^67^. The percentage of aligned reads was above 83% for most samples. Expression count matrices were then processed with an analysis pipeline that used Bioconductor package limma ^68^ for normalization (using the TMM method) and differential expression testing, taking into account only those genes expressed with at least 1 count per million (CPM) in at least two samples (the number of samples for the condition with the least replicates), and using a random variable to define blocks of samples obtained from the same animal. Changes in gene expression were considered significant if associated with a Benjamini and Hochberg adjusted p-value < 0.05. A complementary GSEA ^69^ was performed for each contrast, using the whole collection of genes detected as expressed (12,872 genes) to identify gene sets that had a tendency to be more expressed in either of the conditions being compared. We retrieved gene sets, representing pathways or functional categories from the Hallmark, KEGG, Reactome, and Biocarta databases, and GO collections from the Biological Process, Molecular Function, and Cellular Component ontologies from MsigDB ^70^. Enriched gene sets with a false discovery rate (FDR) < 0.05% were considered of interest.

Data were analyzed with Python 2.7, using the Seaborn (https://seaborn.pydata.org) and Pandas libraries (https://pandas.pydata.org/).

For scRNAseq data processing and in silico EC Selection, the following pipeline was followed. For alignment and quantification of gene expression the reference transcriptome was built using mouse genome GRCm38 and ensembl gene build version 98 (sep2019.archive.ensembl.org). The phiYFP-sv40pA, MbTomato-2A-Cre-WPRE-sv40pa or CreERT2 transgenes sequences expressed in the samples were added to the reference. Gene meta-data were obtained from the corresponding Ensembl BioMart archive. Reads from hashtags and transcripts were processed, aligned, and quantified using the CellRanger v4.0.0. pipeline. The following quality control steps were performed to minimize low-quality cells and improve posterior normalization and analysis: i) A minimum of normalized counts per cell of 1500 and a maximum of 40000 was applied. ii) A minimum gene detection filter of 600 genes. iii) A maximum 25% of MT content. iv) A minimum cell fraction in the sample of 0.1%. The cell fraction refers to the proportion of UMIs the cell contributes with to the total UMIs in the sample. v) Cells with high numbers of reads in just a few genes were filtered out. vi) Cells with more than 0.1% reads in HBB genes were removed. vii) Cells with < 100 UMIs in the hashtags were removed to improve sample calling. Viii) Only single cells were selected; doublets were filtered out.

Cells were de-multiplexed using the pipeline provided with the Seurat package. Counts were normalized and scaled for posterior analysis. Clusters were identified using the nearest neighbors algorithm at different resolutions. Clusters and cells were classified based on the SingleR method using Blueprint Encode and the Human Primary Cell Atlas cell type profiles collection. This identification was used to select ECs for the analysis and remove minor contaminants (T and B cells and monocytes).

### Liver endothelial cell proteomics

#### Preparation of protein extracts and on-filter tryptic digestion

Cells were boiled for 10 min in lysis buffer (4% SDS, 50 mM Tris-HCl, pH 8.5) supplemented with 50 mM iodoacetamide (Sigma-Aldrich) to block reduced Cys residues^71^. After centrifugation at 13,000 rpm, supernatants were collected and protein concentration was determined by the RC/DC Protein Assay (Bio-Rad Laboratories).

Samples were subjected to tryptic digestion using a modification of the filter-aided sample preparation (FASP) technology (Expedeon)^72^. Briefly, the protein extracts (100 μg) were diluted in urea sample solution (USS, 8 M urea in 100 mM Tris-HCl at pH 8.5) and loaded onto the filters. After centrifugation and a washing step with USS, the oxidized Cys residues were reduced with 50 mM DTT (GE Healthcare) in USS for 1 h at room temperature. The samples were then centrifuged, washed with USS, and subsequently alkylated with 50 mM S-methyl methanethiosulfonate (MMTS) (Thermo Fisher Scientific) in USS for 1 h at room temperature. Thereafter, the filters were washed three times with USS and three times with ABC Buffer (50 mM NH_4_HCO_3_ at pH 8.8). Proteins were digested using sequencing grade trypsin (Promega) at a 1:40 (w/w) trypsin:sample ratio at 37°C overnight with gentle agitation.

#### Multiplexed stable isotope labeling of tryptic peptides

The resulting tryptic peptides were mixed with 40 µl of trypsin digestion buffer and recovered by centrifugation at 10,000 rpm for 5 min, after which 50 µl of 500 mM NaCl were added and the filters centrifuged for 15 min at 13,000 rpm. Trifluoroacetic acid was added to a final concentration of 1%, and the peptides were desalted on C18 Oasis HLB extraction cartridges (Waters Corporation, Milford, MA, USA) and dried down. The eluted, cleaned-up peptides were subjected to stable isotope labeling using isobaric tags for relative and absolute quantitation (iTRAQ using 8-plex reagents from AB Sciex). The six differentially tagged samples were then pooled and desalted on Waters Oasis HLB C18 cartridges.

#### Peptide fractionation

For peptide fractionation, the labeled sample was separated into five fractions using the high pH reversed-phase peptide fractionation kit (Thermo Fisher Scientific). Bound peptides were eluted with 1) 12.5% (v/v) ACN; 2) 15% (v/v) ACN; 3) 17.5% (v/v) ACN; 4) 20% (v/v) ACN; and 5) 50% ACN. The fractions obtained were vacuum-dried and stored at −20°C for later use.

#### Liquid chromatography coupled to tandem mass spectrometry

The tryptic peptide mixtures were subjected to nanoscale liquid chromatography coupled to tandem mass spectrometry (LC-MS/MS). High-resolution analysis of samples was performed on an EASY-nLC 1000 liquid chromatograph (Thermo Fisher Scientific) coupled to a Q Exactive HF (Thermo Fisher Scientific) mass spectrometer. Peptides were suspended in 0.1% formic acid, loaded onto a C18 RP nano-precolumn (Acclaim PepMap100, 75 μm internal diameter, 3 μm particle size and 2 cm length, Thermo Fisher Scientific), and separated on an analytical C18 nano-column (EASY-Spray column PepMap RSLC C18, 75 μm internal diameter, 3 μm particle size and 50 cm length, Thermo Fisher Scientific) in a continuous gradient: 8-27% B for 240 min, 31-100% B for 2 min, 100% B for 7 min, 100-2% B for 2 min and 2% B for 30 min (where A is 0.1% formic acid in HPLC-grade water and B is 90% ACN, 0.1% formic acid in HPLC-grade water).

Full MS spectra were acquired over the 400-1500 mass-to-charge (m/z) range with 120,000 resolution, 2 x 10^5^ automatic gain control, and 50 ms maximum injection time. Data-dependent MS/MS acquisition was performed at 5 x 10^4^ automatic gain control and 120 ms injection time, with a 2 Da isolation window and 30 s dynamic exclusion. Higher-energy collisional dissociation of peptides was induced with 31% normalized collision energy and analyzed at 35,000 resolution in the orbitrap.

#### Protein identification based on database searching

Proteins were identified in the raw files using the SEQUEST HT algorithm integrated in Proteome Discoverer 2.1 (Thermo Fisher Scientific). MS/MS scans were matched against the mouse (UniProtKB 2017-07 release) protein database supplemented with pig trypsin and human keratin sequences. Parameters for database searching were as follows: trypsin digestion with maximum 2 missed cleavage sites allowed, precursor mass tolerance of 800 ppm, and a fragment mass tolerance of 0.02 Da. N-terminal and Lys iTRAQ 8-plex modifications were chosen as fixed modifications, whereas Met oxidation, Cys carbamidomethylation, and Cys methylthiolation were chosen as variable modifications. SEQUEST results were analyzed with the probability ratio method^73^. An FDR for peptide identification was calculated based on searching against the corresponding inverted database using the refined method^74,75^ after precursor mass tolerance postfiltering at 20 ppm. Quantitative information was extracted from the intensity of iTRAQ reporter ions in the low-mass region of the MS/MS spectra^76^.

#### Statistical analysis of proteomics data

Comparative analysis of protein abundance changes was conducted with the SanXoT software package^77^, designed for the statistical analysis of high-throughput, quantitative proteomics experiments and based on the Weighted Scan, Peptide and Protein (WSPP) statistical model^78^. As input, WSPP uses a list of quantifications in the form of log2-ratios (each cell sample versus the mean of the three WT cell samples) with their statistical weights. From these, WSPP generates the standardized forms of the original variables by computing the quantitative values expressed in units of standard deviation around the means (Zq). Known artifact proteins (keratin contaminants and trypsin) were excluded from the data sets after the analysis.

For the study of coordinated protein alterations, we used the Systems Biology Triangle (SBT) algorithm, which estimates functional category averages (Zc) from protein values by performing the protein-to-category integration, as described^76^. The protein category database was built up using annotations from the Gene Ontology (GO) database.

## Statistical analysis

Comparisons between two groups of samples with a Gaussian distribution were by unpaired two-tailed Student *t*-test. Comparisons among more than two groups were made by 1 way ANOVA followed by the Turkey pairwise comparison. Graphs represent mean +/- SD as indicated, and differences were considered significant at p < 0.05. All calculations were done in Excel, and final datapoints were analyzed and represented with GraphPad Prism. No randomization or blinding was used, and animals or tissues were selected for analysis based on their genotype, the detected Cre-dependent recombination frequency, and the quality of multiplex immunostaining. Sample sizes were chosen according to the observed statistical variation and published protocols.

## Data availability

The bulk RNA-seq, proteomics, and scRNAseq data can be viewed at the Gene Expression Omnibus (GEO) under accession number XXX. In this study, we used the MsigDB as indicated above. All other data supporting the study findings are available from the corresponding author upon request. This includes additional raw data such as unprocessed original pictures and independent replicates, which are not displayed in the article but are included in the data analysis in the form of graphs. Source data are provided with this paper.

## Acknowledgments

Research in the Benedito laboratory was supported by the European Research Council (ERC) Starting Grants AngioGenesHD (638028) and Consolidator Grant AngioUnrestUHD (101001814), the CNIC Intramural Grant Program Severo Ochoa (11-2016-IGP-SEV-2015-0505), the Ministerio de Ciencia y Innovación (MCIN SAF2013-44329-P, RYC-2013-13209, SAF2017-89299-P) and “la Caixa” Banking Foundation (project code HR19-00120). Jesus Vazquez lab was supported by MCIN (PGC2018-097019-B-I00, PID2021-122348NB-I00) and La Caixa (HR17-00247 and HR22-00253). The CNIC is supported by the Instituto de Salud Carlos III (ISCIII), the Ministerio de Ciencia e Innovación (MCIN) and the Pro CNIC Foundation, and is a Severo Ochoa Center of Excellence (grant CEX2020-001041-S funded by MICIN/AEI/10.13039/501100011033).

Microscopy experiments were performed at Microscopy and Dynamic Image Unit, CNIC, ICTS-ReDib, co-funded by MCIN/AEI /10.13039/501100011033 and FEDER “Una manera de hacer Europa” (#ICTS-2018-04-CNIC-16).

M.F.-C. was supported by PhD fellowships from Fundación La Caixa (CX_E-2015-01) and Boehringer Ingelheim travel grants. S. M. was supported by the Austrian Science Fund (FWF) project J4358. A.R. was supported by The Youth Employment Initiative (YEI) PEJD-2019-PRE/BMD-16990. L.G.-O. was supported by the Spanish Ministry of Economy and Competitiveness (PRE2018-085283).

We thank S. Bartlett for English editing; S. F. Rocha for manuscript comments, editing and model figure design; the members of the CNIC Transgenesis, Microscopy, Genomics, Citometry and Bioinformatic units. We also thank F. Radtke, R. H. Adams, F. Alt, B., T. Honjo, I. Flores, J. Lewis, J. Owen, M. Aguet, T. Gridley for sharing the *Dll4^floxed^, Cdh5(PAC)-creERT2, Myc ^floxed^, Rbpj ^floxed^, the p21−/−, Jag1^floxed^, Dll1^floxed^, Jag2^floxed^ Notch1^floxed^ and Notch2^floxed^* mice respectively.

## Author contributions

M.F.-C. and R.B. designed most of the experiments, interpreted results, assembled figures and wrote the manuscript. S.M., L.G-O and A.R. designed and performed scRNAseq experiments. A.B and A. D. supported genomics and bulk/single cell sequencing experiments. M.F-C, A.R, C.T. and F.S-C. analysed bulk and scRNAseq data. E.C. and J.V. designed, performed and analysed proteomics experiments. M.L and M.S.S.-M gave general technical assistance, managed the mouse colonies and breedings. V.C.-G. gave general technical assistance and established some of the ES cell lines used to generate mouse models. H.C. supported the bulk RNAseq data analysis.

---

**Supplementary Figure 1:**
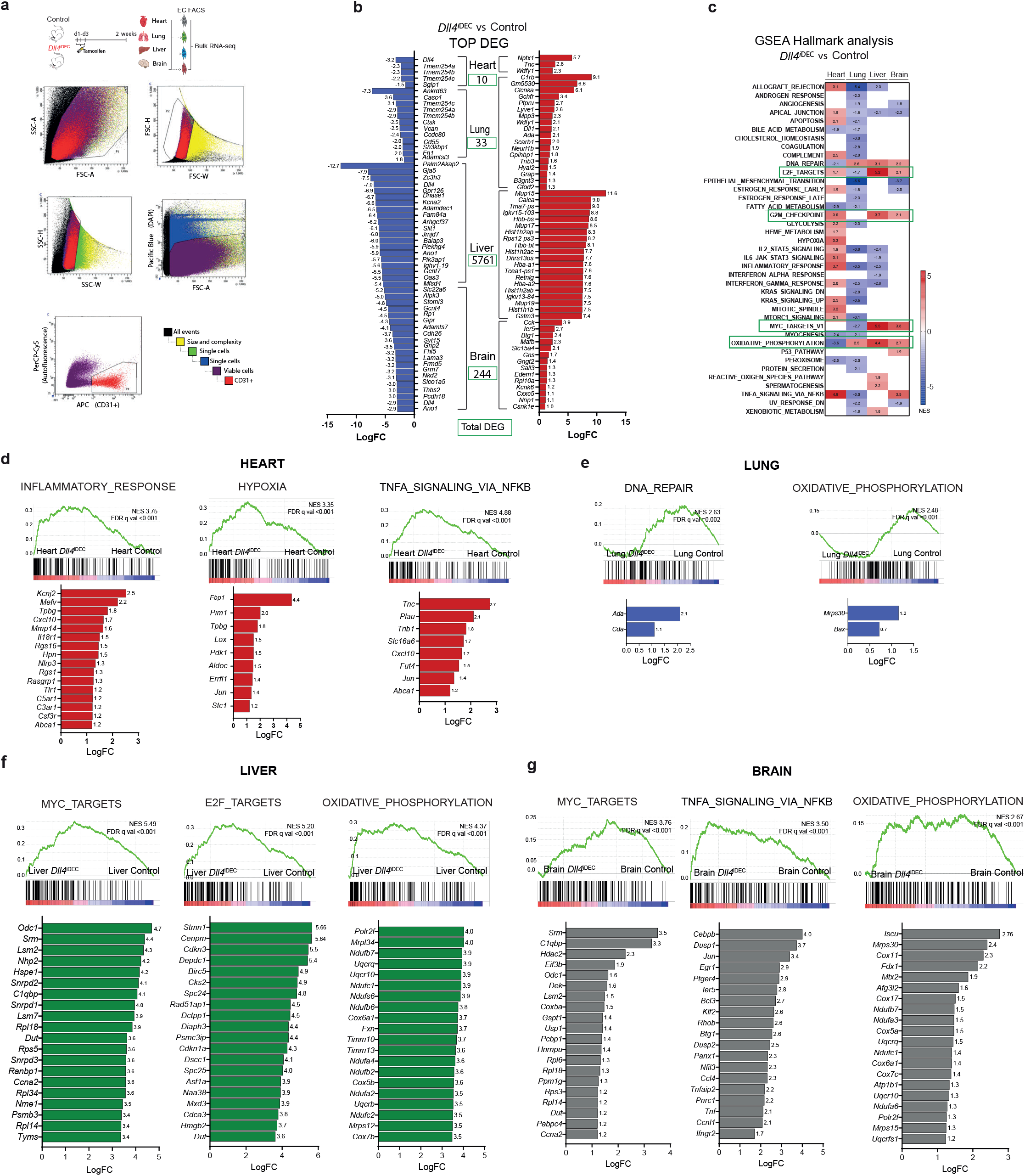
*Dll4* deletion elicits different gene expression signatures among different organ vascular beds. **a,** Schematic representation to illustrate the method used to obtain ECs for bulk RNA-seq analysis. FACS plots show the ECs gating strategy. The detectors, dyes and fluorophores are indicated in the X and Y-axes. **b,** List of the 20 most up- and downregulated genes from the list of differentially expressed genes (DEG, absolute number boxed in green) based on the Benjamini and Hochberg adjusted p-value < 0.05. **c,** Heatmap with the normalized enrichment score (NES) from significant gene set enrichment (GSEA) hallmark analysis (FDR qval< 0.05). **d,** List of upregulated genes in Dll4 mutant ECs within the top 3 enriched gene sets from GSEA hallmark analysis in Heart. **e,** List of upregulated genes within the only enriched gene sets from GSEA Hallmark analysis in Lung *Dll4*^iDEC^ ECs compared to Control ECs. **f,** List of upregulated genes within the top 3 enriched gene sets from GSEA hallmark analysis in Liver *Dll4*^iDEC^ECs compared to Control ECs. **g,** List of upregulated genes within the top 3 enriched gene sets from GSEA Hallmark analysis in Brain *Dll4*^iDEC^ ECs compared to Control ECs. LogFC: Logarithmic Fold Change.

**Supplementary Figure 2:**
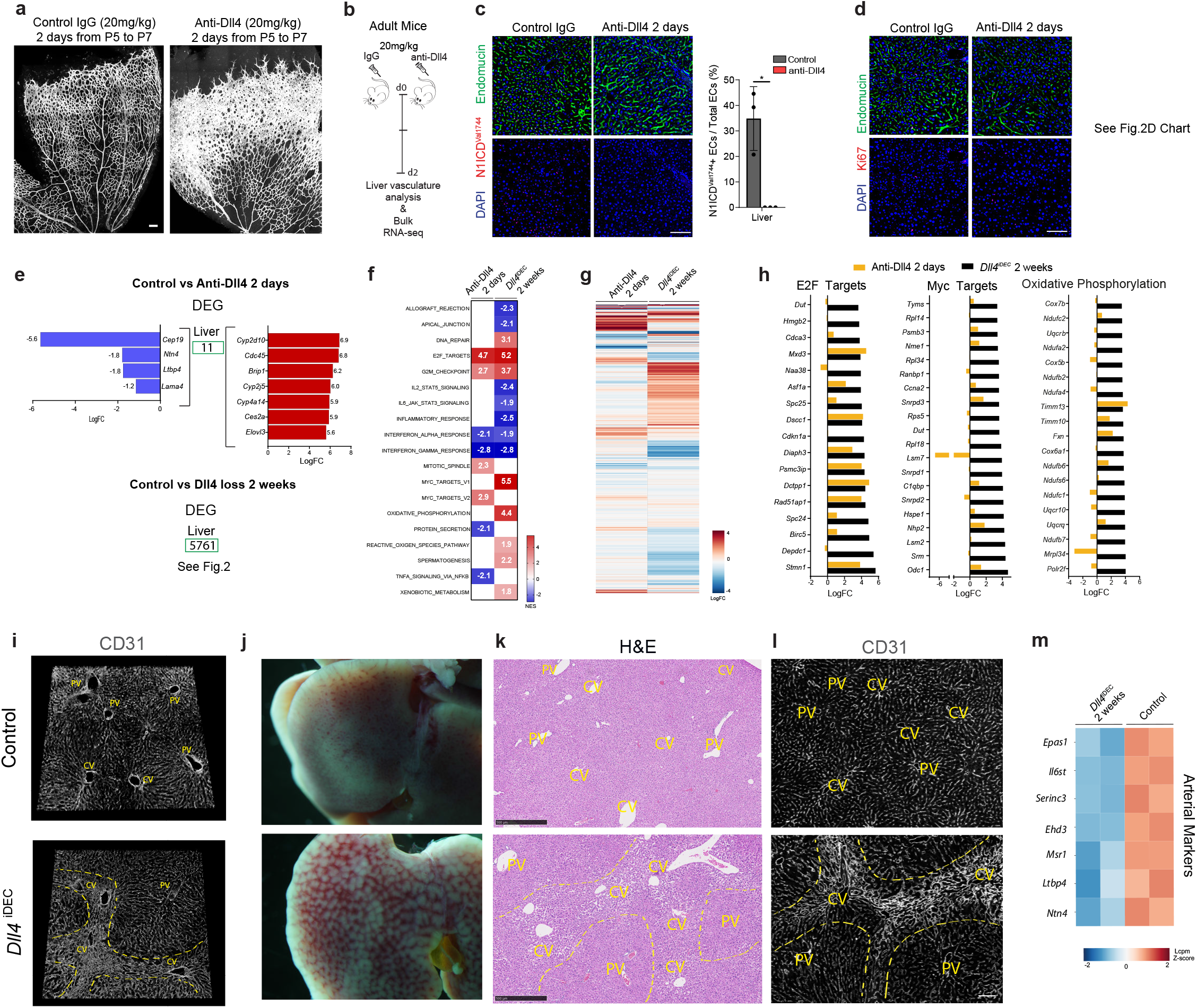
No major genetic and vascular changes after blocking Dll4 signalling in quiescent vessels for 2 days. **a,** Anti-Dll4 treatment for 48h in postnatal day 5 to 7 mice induces a strong increase in retina vascular density and angiogenesis. **b,** Experimental layout for the antibody-based blockade of Dll4 ligand function in adult mice. **c-d,** Confocal micrographs showing that anti-Dll4 blockade for 2 days in adult mice significantly reduces Notch1 activity (cleaved Notch1^Val1744^), but not EC density (DAPI+Endomucin+) and EC proliferation (Ki67+DAPI+ Endomucin+ cells) as depicted in chart D from Figure 2. **e,** List of the few differential expressed genes (DEG) 2 days after Dll4 blockade in liver endothelium. Only 11 genes were differentially expressed, in contrast to the 5761 DEG after targeting Dll4 for 2 weeks. **f,** Heatmap with the normalized enrichment score (NES) from significanty deregulated GSEA hallmark pathways (FDR qval< 0.05). Note that ECs with loss of Dll4 for 2 weeks have more upregulated or downregulated gene sets compared to anti-Dll4 treatment for only 2 days. **g,** Heatmap representing logFC of every expressed gene in the indicated conditions versus control livers. **h,** Comparison of gene expression fold changes between Anti-Dll4 for 2 days (short-term) and Dll4 deletion for 2 weeks (long-term). The top20 DEG belonging to the indicated GSEA pathways are shown. **i,** 3D projection images from confocal scanning of thick vibratome sections show that the vascular malformations observed in *Dll4*^iDEC^ livers are located in sinusoids connecting central veins (CV), but not in sinusoids located close to Portal Veins (PVs). **j,** Low magnification stereomicroscope images of livers from control and *Dll4*^iDEC^ mice showing liver pathology and blood accumulation in the mutants. **k,** Hematoxylin and Eosin staining images of liver sections from control and *Dll4*^iDEC^ mice show sinusoidal dilation in areas surrouding and connecting central veins (CV). **l,** Confocal micrographs showing higher EC density (CD31+) and abnormal or enlarged sinusoids around central veins (delimited by yellow dashed lines). **m,** Heatmap showing that several arterial markers are downregulated in *Dll4*^iDEC^ liver ECs 2 weeks after tamoxifen induction. Charts have n=3 animals per group. Error bars indicate SD. *p < 0.05. Scale bars, 100 μm.

**Supplementary Figure 3:**
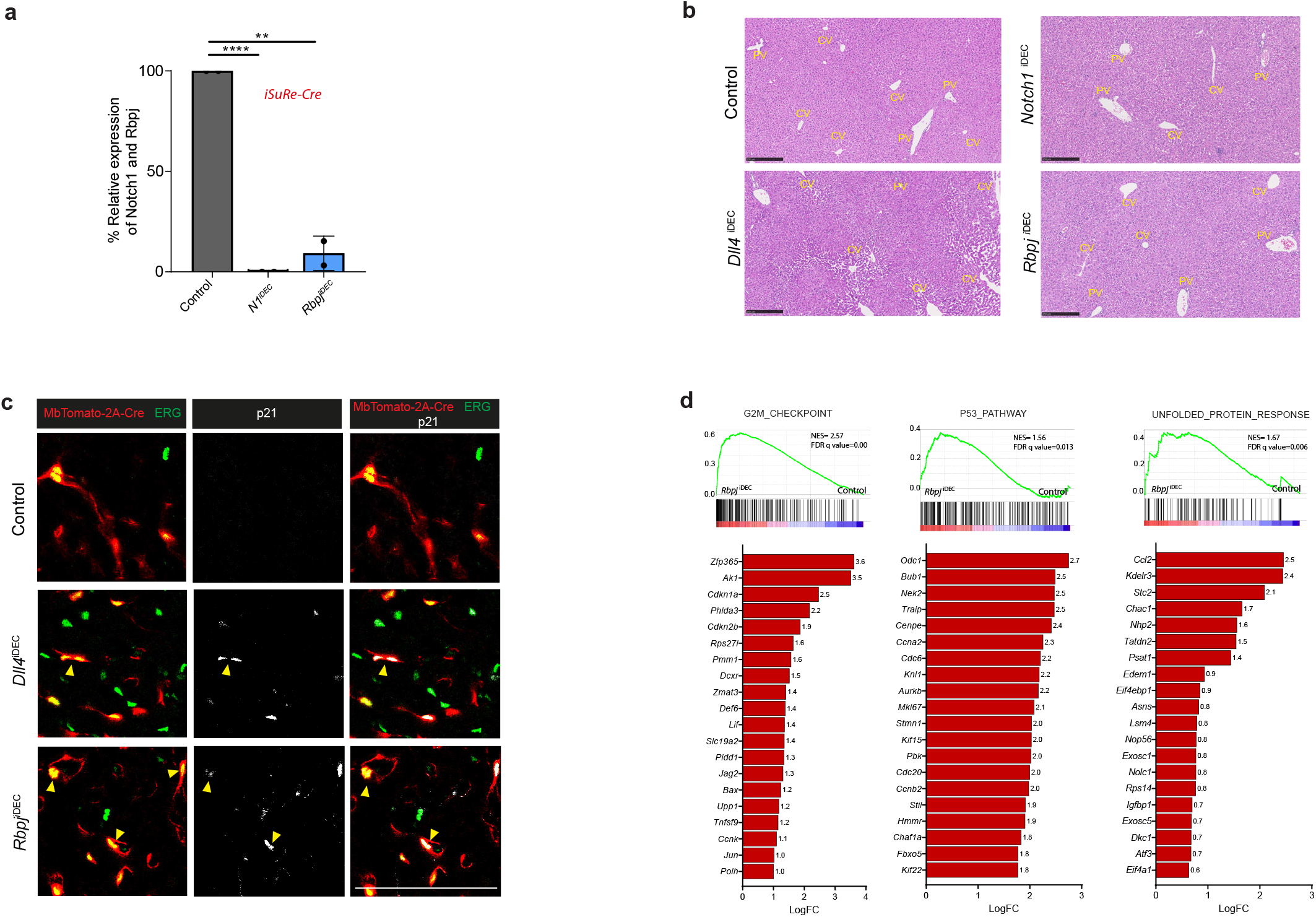
The non-pathologic and arrested endothelial status of *Rbpj* mutant ECs is linked to the upregulation of genes related with the p53, p21, G2M checkpoints, and general replicative or cellular stress genetic pathways. **a,** *Rbpj* and *Notch1* genes were efficiently deleted in liver quiescent ECs as shown by their relative RNA-seq counts per million. Note for *Rbpj gene* only deleted exons reads were quantified. **b,** Representative Hematoxylin and Eosin staining images of liver sections showing strong liver sinusoidal dilation around central veins (CV) and pathology in *Dll4^i^*^DEC^ but not in *Notch1*^iDEC^ or *Rbpj^i^*^DEC^ mutants. Scale bar, 250μm. **c,** Confocal micrographs of liver sections showing that binucleated *Dll4^i^*^DEC^ and *Rbpj^i^*^DEC^ ECs are p21+. Yellow arrowheads indicate p21+ binucleated EC events. **d,** List of the top 20 upregulated genes in Rbpj mutant ECs within the indicated gene sets from the GSEA Hallmark analysis. Charts n=2 animals per group minimum. Error bars indicate SD. **p < 0.01. ****p <0.0001. ns, non-significant. Scale bar, 100 μm.

**Supplementary Figure 4:**
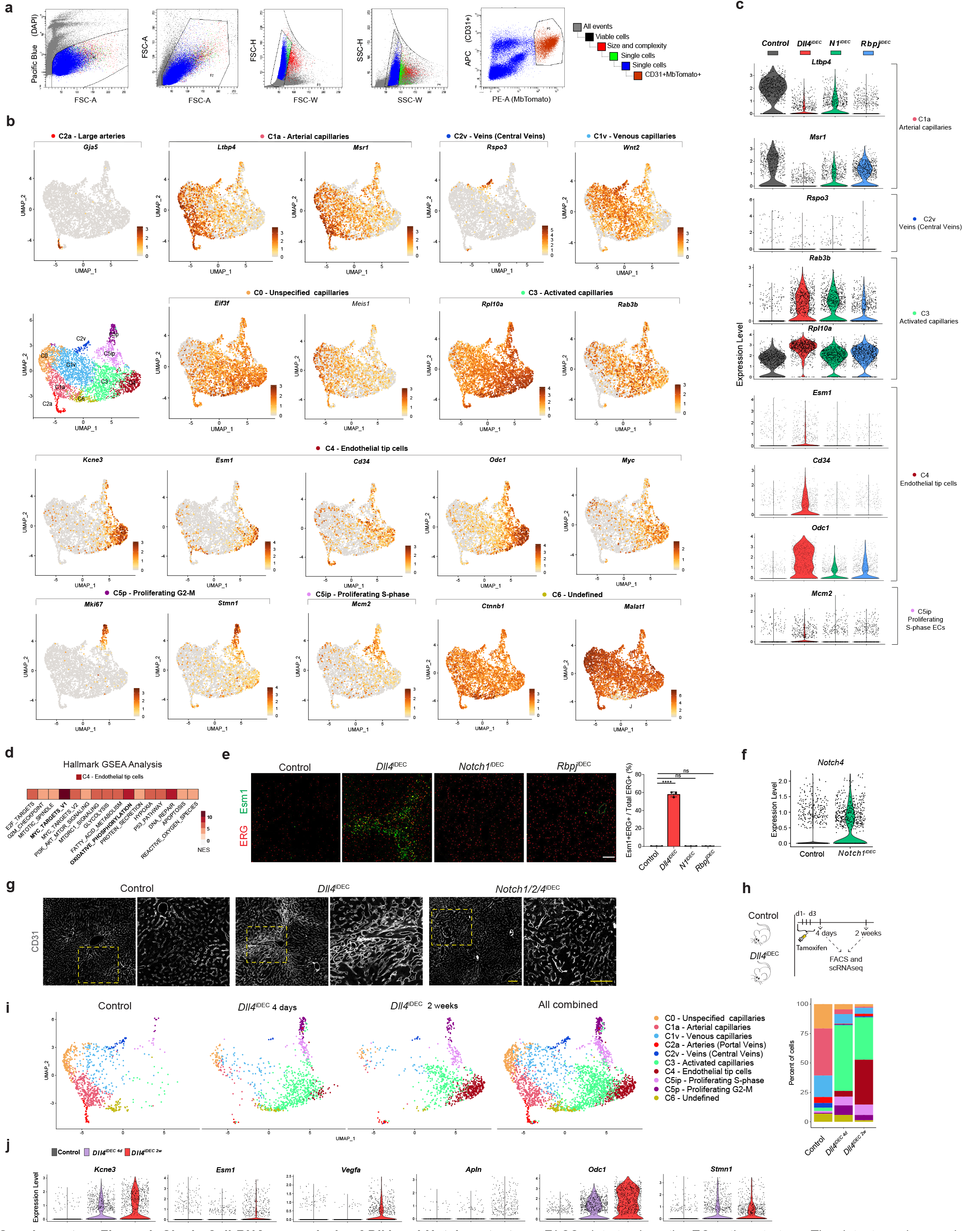
Single Cell RNAseq analysis of Dll4 and Notch mutants. **a,** FACS plots to show the EC gating strategy. The detectors, dyes and fluorophores are indicated in the X and Y-axes. **b,** Feature plots of cluster specific or cluster enriched genes. Some clusters are also characterized by the lack of expression of a given gene. **c,** Violin plots of different cluster markers expression in indicated mutants. **d,** Hallmark GSEA analysis of C4 cluster showing that Myc targets and Oxidative Phosphorylation related genes are the most upregulated pathways. NES. Normalized Enrichment Score. **e,** Confocal micrographs of liver sections showing presence of Esm1+ ECs exclusively in *Dll4*^iDEC^ mutants. **f,** Violin plot showing *Notch4* upregulation in *Notch1*^iDEC^ liver ECs. **g,** Confocal micrographs of liver sections showing abnormal vasculature (CD31+) around central veins (CV) in *Dll4*^iDEC^ livers, but not in *Notch1/2/4*^iDEC^. Yellow dashed rectangle within left panel is to highlight the location of high-magnification images shown in right panel. **h,** Experimental layout for the *Dll4* deletion induction and scRNAseq analysis of *Dll4*^iDEC^ livers. **i,** UMAPs and barplots plot show that full loss of Dll4 signalling for 4 days leads to the loss of the arterial program (C1a) and activation and proliferation of the cells (C3 and C5), but not fully diffferentiated tip cells (C4). **j,** Violin plots showing that targeting Dll4 in quiescent vessels induces a relatively slow and progressive gene expression change in tip-cell related genes. Chart data from n=3 mice. Error bars indicate SD. ****p <0.0001. Scale Bars 100um.

**Supplementary Figure 5:**
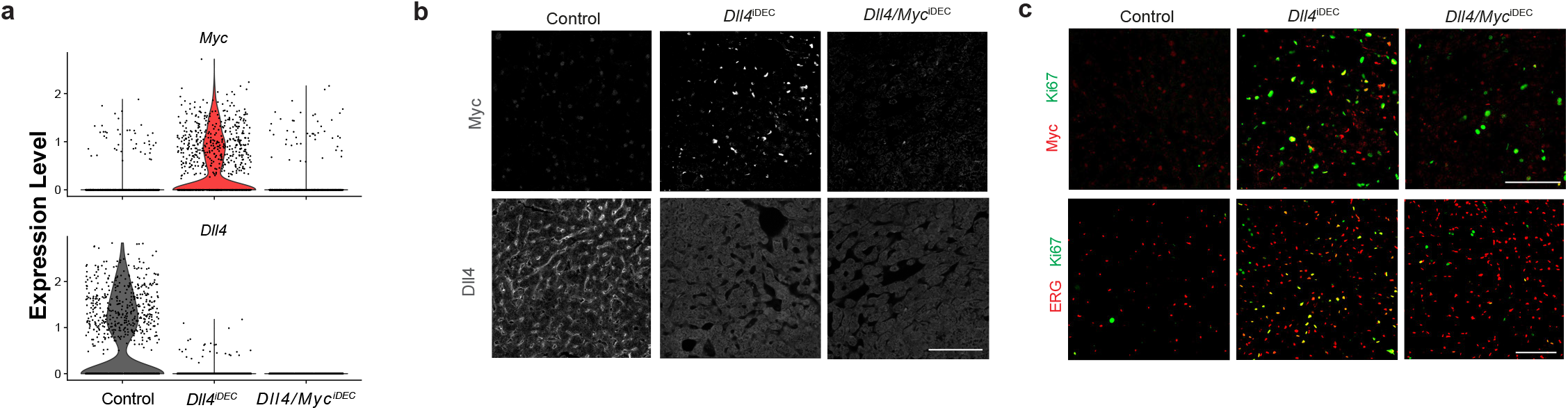
Deletion of *Myc* in *Dll4*^iDEC^ mutants blocks EC proliferation but not vessel enlargement and malfunction. **a,** scRNAseq Violin plots show complete deletion of *Dll4* and *Myc* genes in the indicated samples. **b,** Confocal micrographs of liver sections showing Dll4 absence and Myc upregulation in *Dll4*^iDEC^ livers and absence of Dll4 and Myc in the double *Dll4/Myc*^iDEC^ mutants. **c,** Confocal micrographs of liver sections showing absence of EC proliferation (Ki67+ERG+) and Myc expression in the double *Dll4/Myc*^iDEC^ mutants. Scale bars, 100 μm.

**Supplementary Figure 6:**
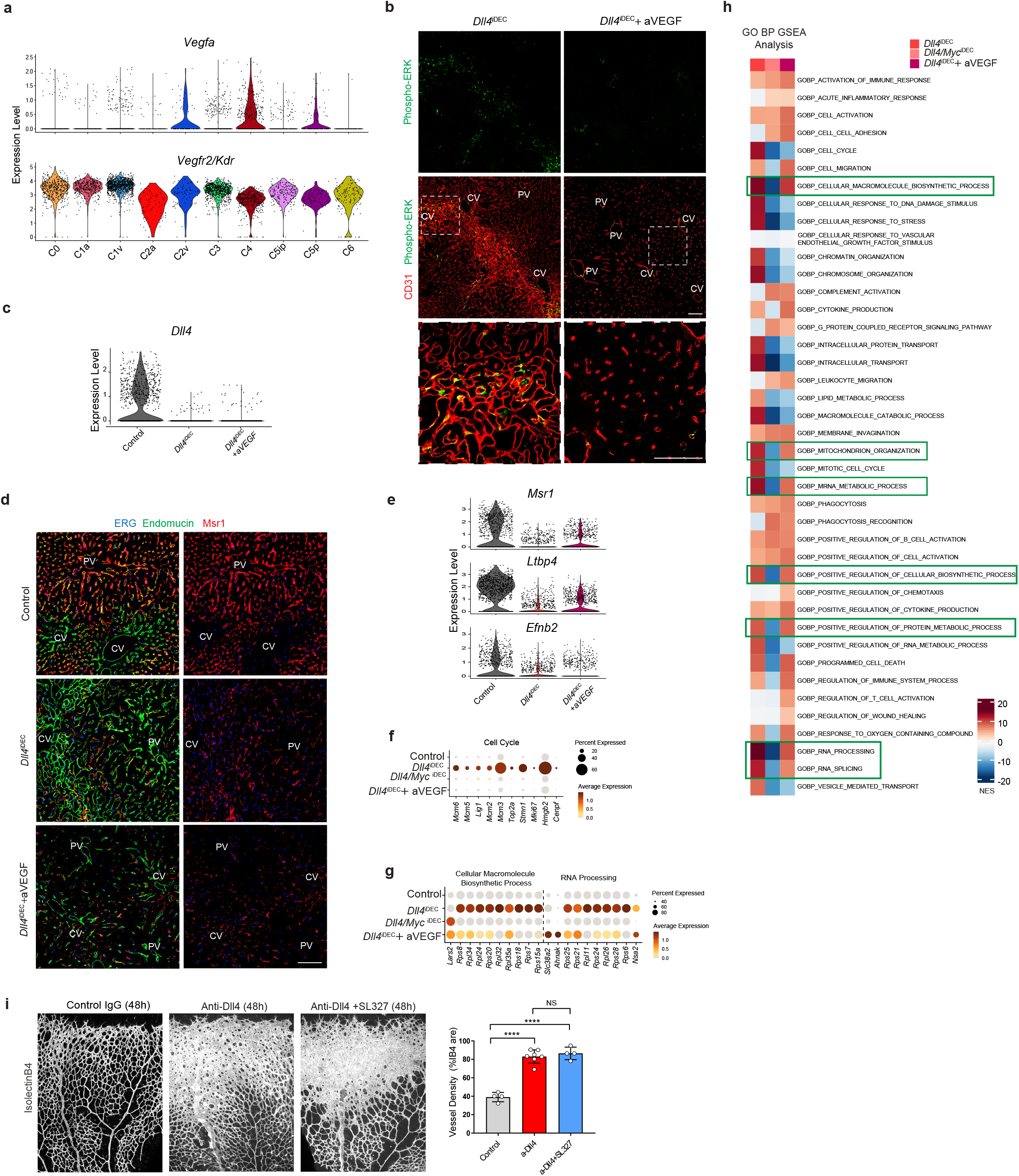
Anti-VEGFA in *Dll4*^iDEC^ mutants reverses EC proliferation, liver pathology and vessel enlargment but not most of the *Dll4*^KO^ genetic programme. **a,** scRNAseq Violin plot showing VEGFA upregulation in the *Dll4*^iDEC^ specific endothelial tip cell cluster (C4) and Kdr/Vegfr2 expression. **b,** Confocal micrographs of liver sections showing the absence of phosphorylation of the VEGFA target ERK after VEGFA blockade. **c,** scRNAseq Violin plot showing *Dll4* deletion in *Dll4*^iDEC^ and *Dll4*^iDEC^+anti-VEGFA samples. **d,** Confocal micrographs of liver sections showing the loss of the capillary marker Msr1 in *Dll4*^iDEC^+anti-VEGFA samples as observed in *Dll4*^iDEC^ liver ECs. **e,** Violin Plots for the arterial markers *Msr1, Ltbp4* and *Efnb2* showing that anti-VEGF does not rescue the arterial identity of cells after *Dll4* deletion. **f,** Dot plot of cell cycle genes showing that ECs are mostly quiescent in *Dll4*^iDEC^+ anti-VEGFA samples. **g,** Dot plot of Cellular Macro-molecule Biosynthetic process and RNA processing GO gene sets showing that they are still active in *Dll4*^iDEC^+ anti-VEGFA samples. **h,** scRNAseq Gene Ontology (GO) analysis of *Dll4*^iDEC^, *Dll4/Myc*^iDEC^ and *Dll4*^iDEC^+anti-VEGFA livers showing that the loss of Myc more strongly downregulates the genes uppregulated in *Dll4*^iDEC^ mutants than the blockade of VEGFA. **i,** SL327 treatment for 48h in pups from postnatal day 5 to 7 does not prevent the increase in vascular density and angiogenesis observed after anti-Dll4. n=4 animals minimum per group. Error bars indicate SD. ****p< 0.0001. ns, non-significant. Scale bars, 100 μm.

**Supplementary Figure 7:**
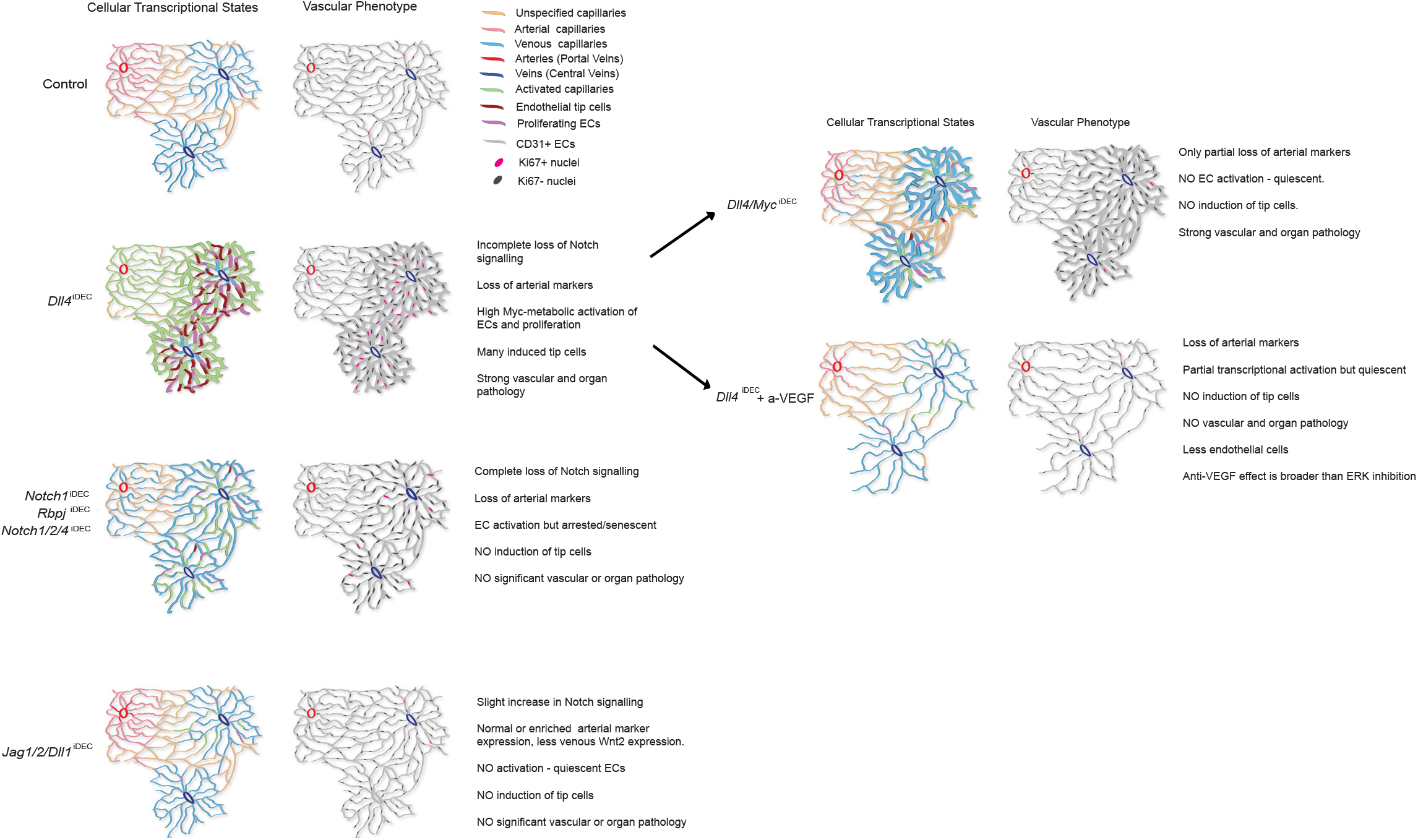
Incongruence between cell states, vascular morphology and pathophysiology. Figure showing the summary of main findings. Targeting of the ligand Dll4 triggers incomplete loss of Notch signalling which results in a strong Myc-driven metabolic activation associated with a well defined cluster of tip cells located around central veins. Dll4 targeted cells have very high ribosome biogenesis, protein synthesis and oxidative phosphorylation favouring cell growth and metabolism. This genetic activation correlates with strong EC proliferation, sprouting, vascular expansion and subsequent organ pathology. Targeting Myc reverses most of the Dll4 mutant genetic activation and cellular states, but Dll4/Myc^iDEC^ mutant livers still have larger sinusoids and organ pathology confirming the incongruence between cell transcriptional and phenotypic states, vascular morphology and organ pathology. Targeting VEGF only partially reduces the Dll4 mutant genetic programs, but it is enough to prevent most of the activated and tip cell states, being the cells in a quiescent state. It also induces the loss of endothelial cells, which overall results in the reversion of the vascular expansion and organ pathology. The effect of Anti-VEGF is broader than and independent of ERK and Myc functions. The loss of Notch receptors or Rbpj leads to complete loss of Notch signalling and also the loss of the arterial transcriptional program, but in this case cells undergo an hypermitogenic MAPK-driven cell-cycle arrest and show cellular senescence features. In contrast to Dll4 mutant cells, these do not effectively proliferate or sprout and only minor vascular and organ pathology is observed. Loss of all other Notch ligands leads to increased Notch signalling, without any associated vascular activation or pathology. This work shows that single cell states often do not correlate with observed vascular phenotypes and organ pathology.

